# Surface anchoring of the *Kingella kingae* galactan is dependent on the lipopolysaccharide O-antigen

**DOI:** 10.1101/2022.06.13.495970

**Authors:** Nina R. Montoya, Eric A. Porsch, Vanessa L. Muñoz, Artur Muszyński, Jiri Vlach, David K. Hahn, Parastoo Azadi, Matthew Sherman, Hyojik Yang, Courtney E. Chandler, Robert K. Ernst, Joseph W. St. Geme

## Abstract

*Kingella kingae* is a leading cause of bone and joint infections and other invasive diseases in young children. A key *K. kingae* virulence determinant is a secreted exopolysaccharide that mediates resistance to serum complement and neutrophils and is required for full pathogenicity. The *K. kingae* exopolysaccharide is a galactofuranose homopolymer called galactan and is encoded by the *pamABC* genes in the *pamABCDE* locus. In this study, we sought to define the mechanism by which galactan is tethered on the bacterial surface, a prerequisite for mediating evasion of host immune mechanisms. We found that the *pamD* and *pamE* genes are glycosyltransferases and are required for synthesis of an atypical lipopolysaccharide (LPS) O-antigen. The LPS O-antigen in turn is required for anchoring of galactan, a novel mechanism for association of an exopolysaccharide with the bacterial surface.

**Significance:** *Kingella kingae* is an emerging pediatric pathogen and produces invasive disease by colonizing the oropharynx, invading the bloodstream, and disseminating to distant sites. This organism produces a uniquely multifunctional exopolysaccharide called galactan that is critical for virulence and promotes intravascular survival by mediating resistance to serum and neutrophils. In this study, we established that at least some galactan is anchored to the bacterial surface via a novel structural interaction with an atypical lipopolysaccharide O-antigen. Additionally, we demonstrated that the atypical O-antigen is synthesized by the *pamD* and *pamE* genes, located downstream of the gene cluster responsible for galactan biosynthesis. This work addresses how the *K. kingae* exopolysaccharide can mediate innate immune resistance and advances understanding of bacterial exopolysaccharides and lipopolysaccharides.

## Introduction

*Kingella kingae* is a gram-negative coccobacillus and is a common member of the oropharyngeal microbiota in children ages 6 months to 4 years, generally as a commensal organism. In recent years, PCR-based diagnostics have revealed that *K. kingae* is also a leading cause of osteoarticular infections and other invasive diseases in young children, highlighting the need for a more detailed understanding of *K. kingae* pathogenicity.^1–3^ To reach sites of disease, *K. kingae* must initially colonize the upper respiratory tract, then breach the respiratory epithelium to gain access to the bloodstream, and then survive in the bloodstream.^3–6^ Intravascular survival is driven by specific bacterial factors, including both a capsular polysaccharide and an exopolysaccharide.^5, 7, 8^ Our prototype strain of *K. kingae,* KK01, expresses an exopolysaccharide that is a galactofuranose homopolymer with the structure [➔5)-β-Gal*f*-(1➔]_n_ and is termed the PAM-galactan.^9, 10^ Bendaoud *et al.* identified a five-gene locus designated *pamABCDE* that is necessary for PAM-galactan production and found that only *pamABC* were required for synthesis of the galactofuranose homopolymer in *Escherichia coli.*^10^ In this system, PAM-galactan could be isolated from whole cell sonicates but not the bacterial surface, suggesting that there may be other genes required for secretion. Initially, PAM-galactan was thought to contain abundant DNA and hence was termed <u>p</u>oly*-*DNA-containing <u>a</u>nti-adhesive <u>m</u>aterial extract (PAM extract). Subsequent work established that the galactofuranose polymer does not contain DNA^9, 10^; thus, in this work we will use the term “galactan” to refer to the [➔5)-β-Gal*f*- (1➔]_n_ homopolymer.

While the functions of bacterial exopolysaccharides have traditionally been attributed to processes such as biofilm formation, our work has identified roles for the *K. kingae* galactan exopolysaccharide in immune evasion. In particular, galactan has been shown to play a crucial role in intravascular survival by inhibiting complement-mediated killing and blocking neutrophil phagocytosis.^5, 8^ These novel functions have prompted us to pursue a more detailed understanding of the mechanism of galactan surface presentation and surface anchoring.

Early characterization of *K. kingae* galactan suggested that it is loosely tethered to the bacterial outer membrane through an unknown mechanism, as it could be readily identified in whole bacteria surface washes with phosphate-buffered saline (PBS) as well as mild acid extracts of pre-washed whole bacteria, a technique used to release the polysaccharide capsule from its lipid anchor in the outer membrane.^9^ An example of exopolysaccharide tethering to the membrane has been observed in *E. coli,* which contains a colanic acid exopolysaccharide that is dynamically converted between a lipopolysaccharide (LPS)-linked form, a secreted form, and a membrane-anchored form in response to metabolic stress.^11, 12^ Similarly, the enterobacterial common antigen (ECA) in *Enterobacteriaceae* is tethered to the membrane by either a lipid moiety or a covalent linkage to the lipid A-core oligosaccharide of the LPS.^13–15^

LPS is a multifunctional glycolipid and is highly abundant in the outer membrane of gram-negative bacteria. The functions of LPS are largely influenced by the structure, and there is significant heterogeneity in LPS composition across bacterial species, reflected both in structural diversity and in the genetic heterogeneity of LPS biosynthetic machinery. LPS is anchored in the membrane by lipid A, which is covalently linked to the core oligosaccharide. In bacteria with a smooth LPS, the core oligosaccharide is further connected to an O-antigen composed of repeating sugar units.^16^ LPS is crucial for membrane integrity and creates a selective permeability barrier, inhibiting the diffusion of small hydrophobic molecules, including detergents and antibiotics.^16–18^ Similarly, the length of the O-antigen and the array of covalent LPS modifications serve to restrict access of antibodies and complement proteins to the bacterial surface.^19–22^ In addition, lipid A, which is recognized by Toll-like receptor 4 (TLR4), is a primary driver of the host inflammatory response.^23^ Given the interface between LPS and the host immune system, LPS remodeling is often associated with immune evasion.^24^

In this work, we identify homology between the *pamD* and *pamE* predicted products and LPS-modifying glycosyltransferases. Consistent with this homology, we demonstrate that the *pamD* and *pamE* gene products mediate synthesis of an atypical LPS O-antigen and are essential for surface anchoring of galactan. In addition, we provide strong evidence that galactan is attached to the LPS O-antigen, facilitating galactan-mediated evasion of immune mechanisms.

## Results

### Surface extracts from *pamDE* mutants lack galactan material

To assess the presence of galactan on the bacterial surface, we isolated surface washes from strains KK01 Δ*csaA* (lacking the capsule synthesis locus), KK01 Δ*csaApamABCDE* (a mutant lacking both the capsule synthesis locus and the full *pam* locus), KK01 Δ*csaApamABC* (a mutant lacking the capsule synthesis locus and the *pamABC* genes), and KK01 Δ*csaApamDE* (a mutant lacking the capsule synthesis locus and the *pamDE* genes). Strain KK01 Δ*csaA* was used as the parent strain for this analysis to avoid contamination by the polysaccharide capsule. Surface washes were treated with DNase I, RNase A, and Proteinase K and were then separated on a DOC-PAGE gel and stained with silver. As shown in Figure 1A, the surface wash from strain KK01 Δ*csaA* contained a prominent, broad high molecular weight (HMW) silver-stained band. This band was absent in surface washes from strains KK01 Δ*csaApamABCDE,* KK01 Δ*csaApamABC,* and KK01 Δ*csaApamDE.* Based on previous work demonstrating that deletion of the *pamABCDE* genes results in the loss of galactan,^9^ we wondered whether the HMW band was galactan. To address this possibility, we generated a polyclonal antiserum against purified galactan recovered from the surface of KK01 Δ*csaA* [purity 92.9% galactose as determined by gas chromatography-mass spectrometry (GC-MS); Supplemental Table 1]. This antiserum (GP-19) was adsorbed with an acetone powder of strain KK01 Δ*csaApamABCDE* to remove nonspecific antibodies and was found to be reactive with whole cell sonicates of *E. coli* JM109 harboring *pamABC* on a plasmid but not with whole cell sonicates of *E. coli* JM109 harboring empty vector, confirming reactivity with galactan (Supplemental Figure 1).

**Figure 1:**
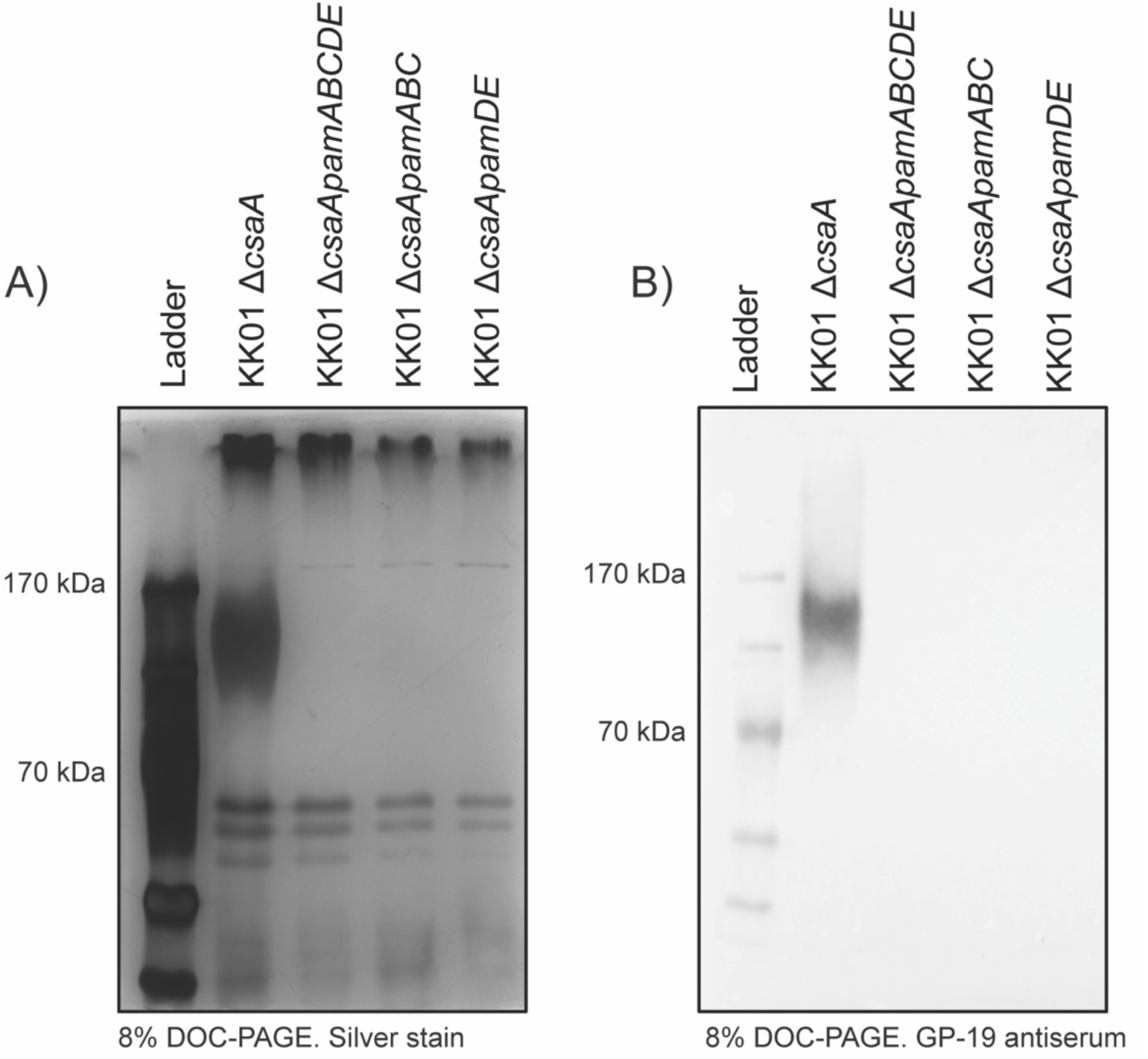
Galactan is not present in surface washes of Δ*csaApamDE* mutants. A) Surface washes were isolated from *K. kingae* strains KK01 Δ*csaA,* KK01 Δ*csaApamABCDE,* KK01 Δ*csaApamABC,* and KK01 Δ*csaApamDE* by vortexing whole bacteria in PBS and concentrating the resulting material. Galactan in the samples was visualized by DOC-PAGE and silver stain. B) The presence of galactan in bacterial surface washes was confirmed with detection by GP-19 antiserum, specific for the *K. kingae* galactan. Representative images are shown for each.

Consistent with these results, Western immunoblots of surface washes performed with the GP-19 antiserum revealed reactivity with strain KK01 Δ*csaA* in the region corresponding to the size of the silver-stained material but no reactivity with strains KK01 Δ*csaApamABCDE,* KK01 Δ*csaApamABC,* or KK01 Δ*csaApamDE* (Figure 1B), indicating that only strain KK01 Δ*csaA* contains galactan on the surface. These results demonstrate that the *pamD* and *pamE* genes are required for surface-associated galactan.

### Wild-type *K. kingae* produces distinctive LPS molecules that migrate in a ladder pattern and as a high molecular weight species and are lost in mutants lacking *pamD* and *pamE*

Homology analysis revealed that the *pamD* and *pamE* gene products share significant homology with GTB-superfamily glycosyltransferases and family 25 LPS modifying glycosyltransferases, respectively. To explore the possible role of the *pam* genes in biosynthesis of the *K. kingae* LPS, we compared the migration patterns of LPS samples prepared from strains KK01 Δ*csaA*, KK01 Δ*csaApamABCDE*, KK01 Δ*csaApamABC*, and KK01 Δ*csaApamDE* using a hot-phenol based method, separated on a DOC-PAGE gel, and stained with silver. As shown in Figure 2A, strain KK01 Δ*csaA* produces LPS that migrates as two distinct modal clusters, including a high molecular weight species at the top of the gel (designated HMW LPS, yellow arrow) and a low molecular weight ladder of distinct bands (designated LMW LPS, white arrow and bracket). The LMW LPS was distinct from the lipooligosaccharide (LOS) produced by *Haemophilus influenz*ae and was notably larger than the LOS truncated after the second heptose residue produced by an *H. influenzae* Δ*rfaF* mutant.^25^ Although distinct from a typical O-antigen produced by *Salmonella enterica* serovar Enterica, the observed ladder pattern suggests that the KK01 Δ*csaA* LMW LPS contains a repeating unit reminiscent of an O-antigen. The KK01 Δ*csaApamABC* mutant lacked HMW LPS, and the KK01 Δ*csaApamDE* mutant lacked both LMW LPS and HMW LPS and displayed only a truncated LPS species, suggesting that the *pamD* and *pamE* genes are required for production of a repeating sugar unit. Of note, the KK01 Δ*csaApamDE* LPS is larger than the LOS produced by a KK01Δ*csaApamABCrfaF* mutant. Homology analysis suggests that the *K. kingae rfaF* gene encodes a lipooligosaccharide heptosyltransferase II, responsible for adding the second heptose residue onto the growing LOS. Deletion of the *K. kingae rfaF* gene resulted in a low molecular weight LOS molecule similar in size to the LOS produced by *H. influenzae* Δ*rfaF.* The KK01 Δ*csaApamABCrfaF* LOS was smaller than the truncated LOS produced by KK01 Δ*csaApamDE*, suggesting that KK01 Δ*csaApamDE* retains residues of the core oligosaccharide.

**Figure 2:**
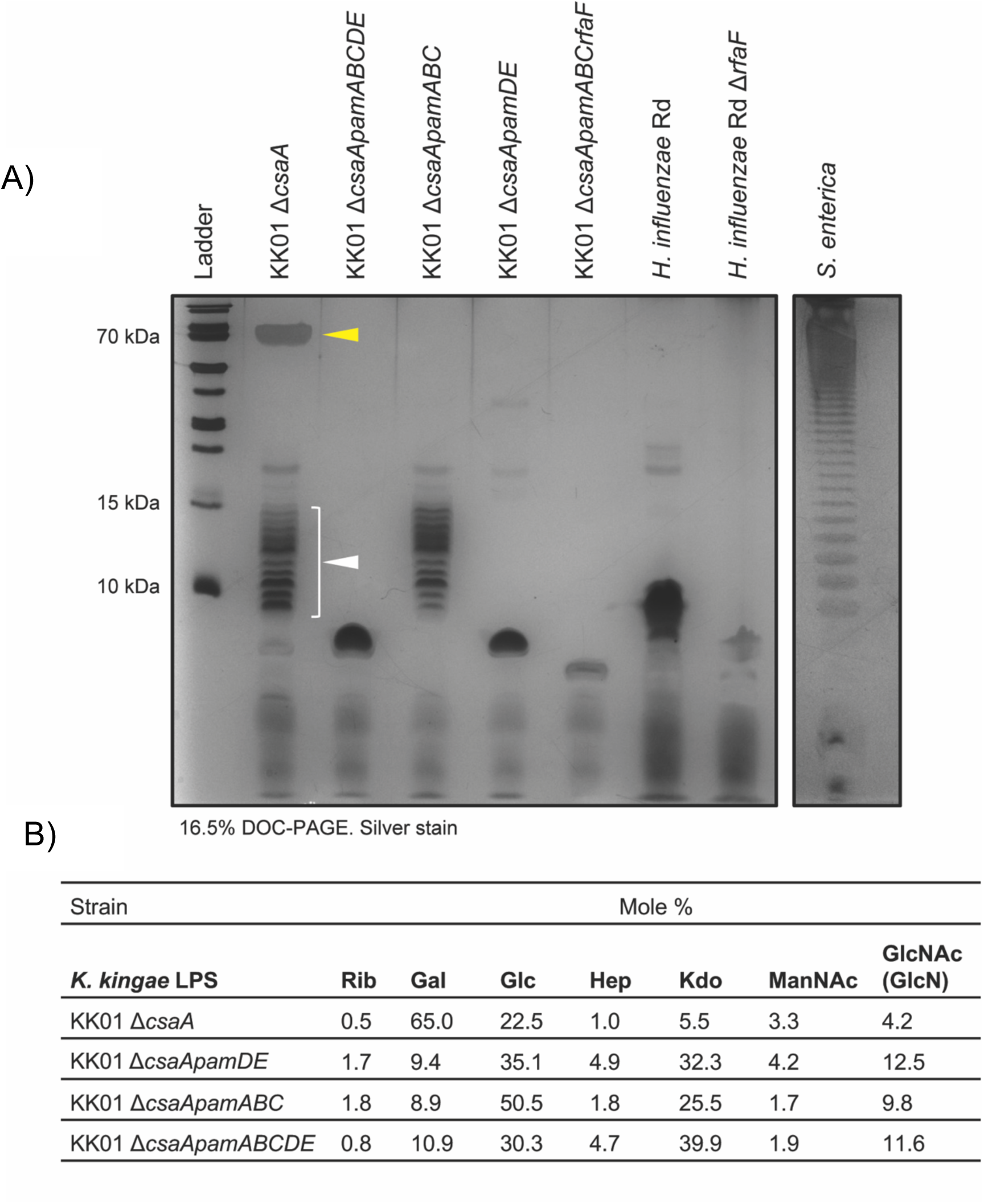
WT *K. kingae* LPS is enriched in galactose and displays two modal clusters of LPS species that are selectively lost in Δ*csaApamABC* and Δ*csaApamDE* mutants. A) LPS was isolated from *K. kingae* strains KK01 Δ*csaA,* KK01 Δ*csaApamABCDE,* KK01 Δ*csaApamABC,* KK01 Δ*csaApamDE,* and KK01 Δ*csaApamABCrfaF* using a hot-phenol extraction protocol. Rough LPS from strains *H. influenzae* Rd and *H. influenzae* Rd Δ*rfaF* and smooth LPS from *S. enterica* were run as controls. The KK01 Δ*csaA* LPS contains two modal clusters of LPS species, the low molecular weight (LMW) LPS indicated by the white arrow and the high molecular weight (HMW) LPS indicated by the yellow arrow. Representative image shown. B) Comparative glycosyl composition analysis of LPS purifed from the *K. kingae* KK01 Δ*csaA* parent strain and the KK01 Δ*csaApamDE,* KK01 Δ*csaApamABC,* and KK01 Δ*csaApamABCDE* mutants. Rib, Ribose; Gal, Galactose; Glc, Glucose; Hep, Heptose; Kdo, 3-deoxy-D-*manno*-octulosonic acid; ManNAc, N-acetylmannosamine, GlcNAc, N-acetylglucosamine ;* the GlcN, glucosamine present in lipid A was converted to GlcNAc during the re-*N*-acetylation step of the chemical derivatization

These results demonstrate that *K. kingae* produces two modal clusters of LPS species: 1) LMW LPS composed of lipid A, core oligosaccharide, and an atypical O-antigen and 2) HMW LPS composed of lipid A, core oligosaccharide, an atypical O-antigen, and an additional large polysaccharide component that may be attached to the O-antigen. These results also establish that the *pamDE* genes are required for production of LMW LPS and that the complete *pamABCDE* locus is required for production of HMW LPS.

### The *K. kingae* LPS is primarily composed of galactose

To determine the carbohydrate composition of the LPS molecules produced by KK01 Δ*csaA* and mutants lacking specific *pam* genes, we performed glycosyl composition analysis of purified LPS samples from these strains. This analysis revealed that galactose was the most abundant monosaccharide in KK01 Δ*csaA* LPS (Figure 2B). In strains KK01 Δ*csaApamABCDE,* KK01 Δ*csaApamABC,* and KK01 Δ*csaApamDE,* the amount of galactose was markedly reduced, while the relative amounts of glucose, heptose, and 3-deoxy-D-manno-octulosonic acid (Kdo) were increased, likely reflecting a shortening of the polysaccharide attached to lipid A. The compositional analysis was consistent with the loss of galactose from HMW LPS in KK01 Δ*csaApamABC* and the loss of both LMW LPS and HMW LPS in strains KK01 Δ*csaApamABCDE* and KK01 Δ*csaApamDE.* These results establish that *K. kingae* HMW LPS contains a large galactose polysaccharide, dependent on all the *pam* genes. Given our understanding of the role of *pamABC* in the biosynthesis of galactan, these data suggest that the presence of galactose in HMW LPS may result from anchoring of galactan to the LMW LPS atypical O-antigen.

### HMW LPS species react with anti-galactan antiserum

To determine if the galactose component of HMW LPS was galactan, we used the GP-19 antiserum to detect galactan material in purified LPS samples from strains KK01 Δ*csaA*, KK01 Δ*csaApamABCDE,* KK01 Δ*csaApamABC,* and KK01 Δ*csaApamDE*. LPS samples were resolved on a DOC-PAGE gel and examined by Western immunoblot with GP-19. As shown in Figure 3, reactivity was observed with LPS from strain KK01 Δ*csaA* but not from strain KK01 Δ*csaApamABCDE* or strain KK01 Δ*csaApamABC*, indicating the presence of galactan in HMW LPS and highlighting the essential role of the *pamABC* genes in galactan biosynthesis. Reactivity was also lacking with LPS from strain KK01 Δ*csaApamDE,* consistent with the conclusion that the *pamD* and *pamE* genes are required for the presence of galactan in HMW LPS. In parallel experiments, we coated ELISA plates with purified LPS samples and examined these plates with the GP-19 antiserum. Like the results observed by Western blot, measurable reactivity was only detected with LPS from strain KK01 Δ*csaA* (Figure 3B). These results indicate that galactan co-purifies with HMW LPS.

**Figure 3:**
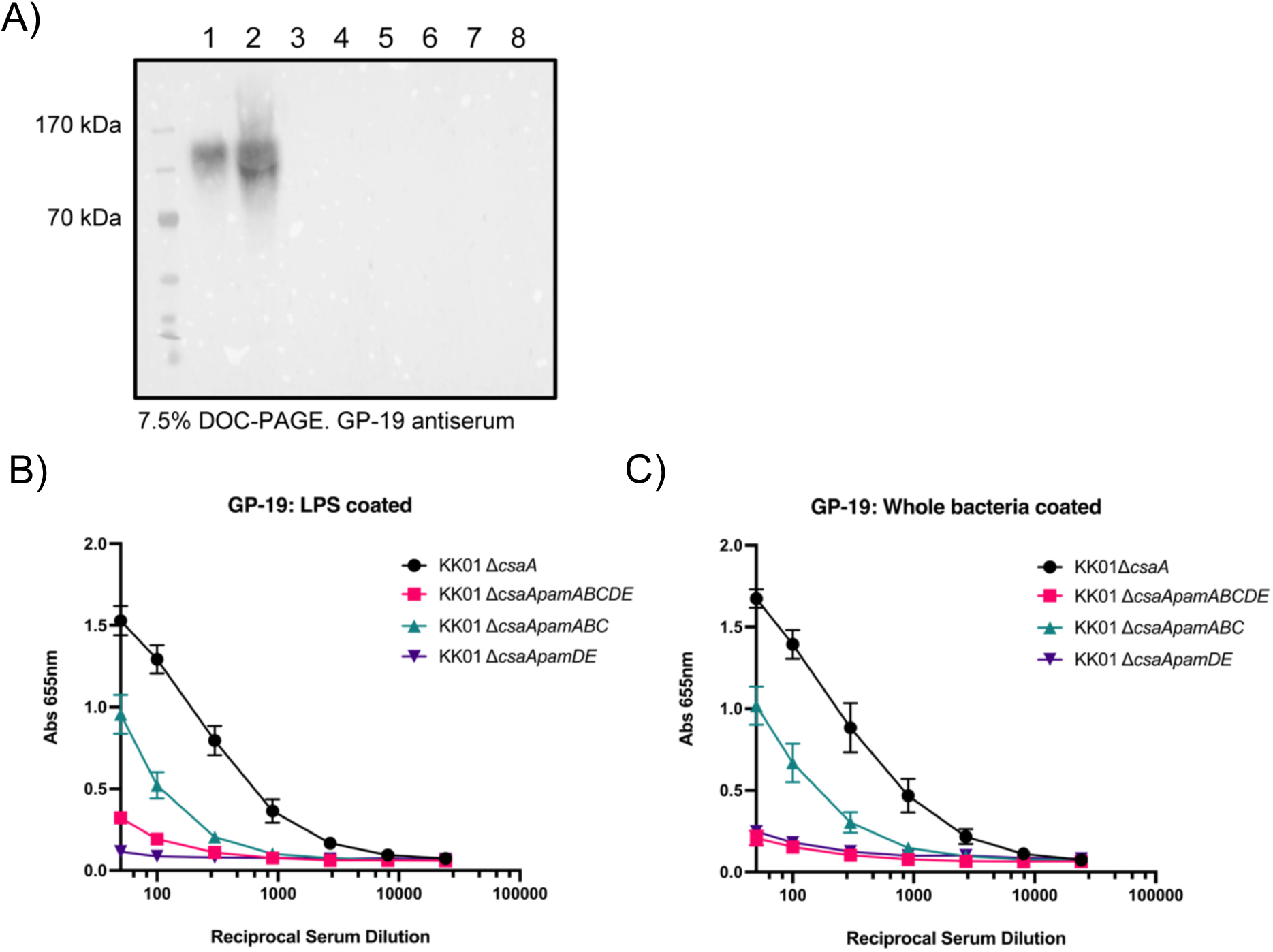
LPS from Δ*pamDE* mutants does not react with GP-19 galactan antiserum. A) LPS samples (odd lanes) and surface washes (even lanes) from *K. kingae* strains KK01 Δ*csaA* (lane 1,2), Δ*csaApamABCDE* (lane 3,4), Δ*csaApamABC* (lane 5,6), and Δ*csaApamDE* (lane 7,8) were separated by DOC-PAGE and galactan was detected with GP-19 antiserum. Representative image shown. B,C) Galactan was detected in purified LPS samples (B) and in whole bacteria (C) with GP-19 antiserum by ELISA. Three biological replicates were performed. Data are expressed as mean ± SEM from three independent experiments.

In addition, the ELISA data revealed that plates coated with LPS from strain KK01 Δ*csaApamABC* generated an intermediate level of reactivity with GP-19 antiserum (Figure 3B). Because reactivity was negligible in plates coated with LPS from strain KK01 Δ*csaApamDE* (Figure 3B), we posited that the intermediate reactivity observed with LPS from KK01 Δ*csaApamABC* was specifically generated against epitopes in LMW LPS produced by the *pamD* and *pam*E gene products, suggesting that the galactan sample used for antibody production also contained LPS. Similar trends were observed when ELISA plates were coated with whole bacteria (Figure 3C).

### HMW LPS can be separated from LMW LPS by size exclusion and is enriched in galactofuranose

To establish definitively that the galactose component of *K. kingae* HMW LPS is identical to galactan ([➔5)-β-Gal*f*-(1➔]_n_), we separated the HMW LPS species from the LMW LPS ladder using size exclusion chromatography (SEC) in the dissociative condition in the presence of a deoxycholic acid detergent. The SEC yielded two significant fractions corresponding to HMW LPS and LMW LPS (Fr 2 and Fr 3, respectively), confirmed by analysis of the individual subfractions [Fr 2 (22-23) and Fr 3 (28-33)] via DOC-PAGE (Figure 4A). A comparative DOC-PAGE analysis of the pooled HMW LPS (Fr 2) and LMW LPS (Fr 3) with the "Whole" unseparated LPS demonstrated successful isolation of HMW LPS from LMW LPS (Figure 4B, compare lanes "Fr 2", "Fr 3" and "Whole"). Rough LPS from *E. coli* Ra and smooth LPS from *S. enterica* were used as controls.

**Figure 4:**
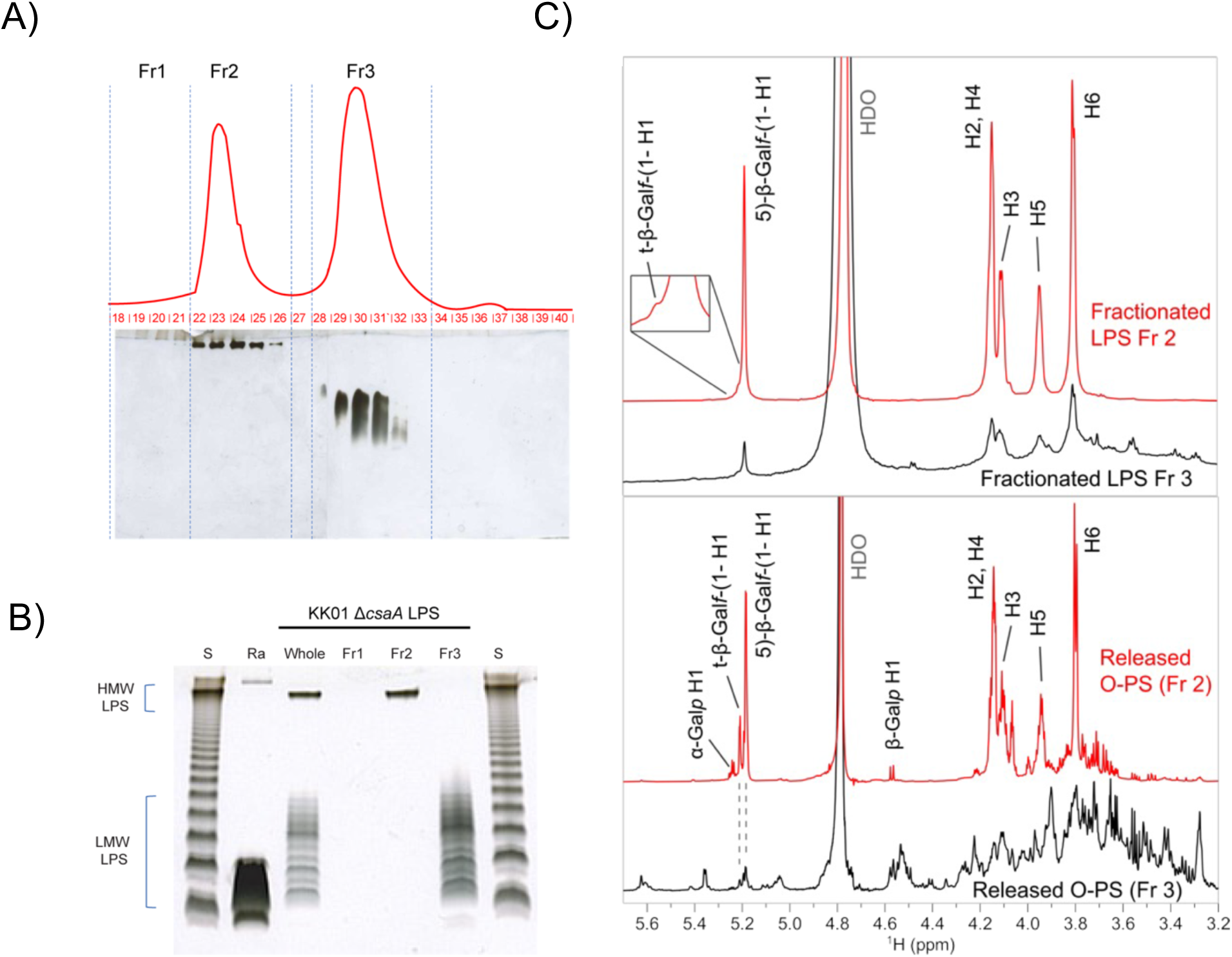
Gal*f* residues are predominantly found in HMW LPS species. A) Size exclusion chromatography of *K. kingae* KK01 Δ*csaA* intact (whole) LPS. The LPS was resolved on the Superdex 75 column, in dissociative conditions, in the presence of deoxycholate. Fractions 1-3 were pooled based on the refractive index (RI) response (red line) combined with the electrophoretic profile of all subfractions (19-40), resolved by DOC-PAGE and silver stained. B) Comparative DOC-PAGE analysis of KK01 Δ*csaA* intact LPS (Whole); and the same LPS fractionated on Superdex 75 (Fr 1-3). Lane S-*Salmonella enterica* serovar Minnesota (S-strain) LPS ; Ra- *E.coli* EH100 LPS (Ra mutant) were used as controls. Each lane was loaded with 1µg of the sample. C) ^1^H NMR spectra of the fractionated LPS (Fr 2 and Fr 3) and the corresponding released O-PS material. The signals of the major Gal*f* polymer are labeled; for simplicity, the ring signals of 5)-β-Gal*f*-(1➔ are marked only as H2–H6. The released O-PS Fr 2 contained free Gal*p* that likely formed from the acid-labile Gal*f* during LPS hydrolysis.

The enriched HMW LPS and LMW LPS fractions (Fr 2 and Fr 3, respectively), as well as the total polysaccharide portion of LPS (O-PS), including core oligosaccharide, O-antigen, and galactan released from each fraction by mild acid hydrolysis, were then analyzed by NMR spectroscopy. The ^1^H spectrum of HMW LPS (Fr 2) contained dominant signals that corresponded to the ➔5)-β-Gal*f*-(1➔ galactan polymer (Figure 4C, top). The identity of the signals was confirmed by an HSQC fingerprint (Supplemental Figure 2) that matched with the data reported previously for galactan^9^ as well as our data obtained on the released O-PS material (see below). The broadened signal appearance and low-intensity signal of the terminal β-Gal*f*-(1➔ group were consistent with a large molecular weight of the polymer. The ^1^H spectrum of LMW LPS (Fr 3) also contained predominantly the same carbohydrate signals as the HMW LPS spectrum (Figure 4C, top), confirming the presence of galactan. The broader appearance of the LMW LPS galactan signals is likely due to the formation of micelles, more pronounced in LMW LPS compared to HMW LPS. ^1^H NMR spectra of the polysaccharides released by acidic hydrolysis from the fractionated LPS are shown in Figure 4C, bottom. The spectrum of the O-PS released from the HMW LPS contained predominantly signals of the galactan that was identified using 2D NMR spectroscopy (Supplemental Figure 3 and Supplemental Table 2). Compared to intact LPS, the released O-PS contained a significantly higher proportion of the terminal β-Gal*f*-(1➔ group, suggesting reduced molecular weight of the polymer. LPS hydrolysis likely resulted in partial degradation of the highly acid-labile Gal*f* polymer, consistent with the presence of a small amount of free Gal*p* (Figure 4C, bottom). The spectrum of the O-PS released from the LMW LPS contained signals of galactan as well as numerous additional signals likely due to O-antigen and core oligosaccharide residues. Overall, the NMR results support the conclusion that the *K. kingae* HMW LPS contains a large ➔5)-β-Gal*f*-(1➔ galactan polymer tethered to the LMW LPS glycolipid region. The LMW LPS contains a relatively small amount of short galactan polymers. This analysis did not allow us to identify a direct covalent linkage between the LMW LPS and galactan, which was likely influenced by the susceptibility of this glycosidic linkage to acid hydrolysis.

To determine if the KK01 Δ*csaApamABC* LPS contains any Gal*f* signals, we performed a comparative NMR analysis. The profile of the anomeric signals in ^1^H NMR spectrum of the total, unfractionated O-PS material released from KK01 Δ*csaApamABC* LMW LPS was very similar to that seen in the KK01 Δ*csaA* LMW O-PS Fr 3 (Supplemental Figure 4A), suggesting similar core and O-antigen structures. Using HSQC, we confirmed that the characteristic Gal*f* signals seen in the KK01 Δ*csaA* LMW O-PS were absent from the spectrum of the Δ*csaApamABC* O-PS (Supplemental Figure 4B), suggesting that the galactose detected in its LPS (Table 1) is in the pyranose form. Given the similarity of the LMW LPS species from KK01 Δ*csaA* and KK01 Δ*csaApamABC,* it is important to note that in the absence of the *pamABC* genes, the LMW LPS is devoid of Gal*f*. Combined, the NMR and the immunochemical analyses show that the galactan polymer is not part of the KK01 Δ*csaApamABC* LPS and that *pamABC* are essential for incorporation of Gal*f* into the O-PS portion of LPS.

### HMW LPS and LMW LPS species are anchored to the same lipid A molecule

In the absence of direct identification of a covalent linkage between LMW LPS and galactan, we sought to determine whether HMW LPS and LMW LPS were both anchored to the same lipid A molecule. Matrix-assisted laser desorption/ionization-time of flight mass spectrometry (MALDI-TOF MS) analysis of the lipid A released from HMW LPS and LMW LPS after hard acid hydrolysis showed that both fractions share identical MS lipid A profiles [Figure 5A, Fr 2 (top spectrum) vs. Fr 3 (lower spectrum)]. A major detected [M-H]^-^ ion at *m/z* 1632.1 could be attributed to a *mono-*phosphoryl-hexa-acyl lipid A species consisting of (GlcN)_2,_P,(14:0(3-OH))_2_,(12:0(3-OH))_2_,(12:0)_2_. We also detected a minor ion at *m/z* 1712.1 representing *bis*-phosphoryl-hexa-acyl lipid A consisting of (GlcN)_2_,P_2_,(14:0(3-OH))_2_,(12:0(3-OH))_2_,(12:0)_2_. The low-intensity ion at *m/z* 1755.1 could be attributed to the *mono*-phosphoryl-hexa-acyl lipid A substituted with phosphoethanolamine (PEtN) and containing (GlcN)_2_,P,PEtN (14:0(3-OH))_2_,(12:0(3-OH))_2_,(12:0)_2_. The relatively lower intensities of [M-H]^-^ ions corresponding to the *bis*-phosphoryl or PEtN-substituted lipid A in the MS analysis were likely caused by a susceptibility of these groups to 1% acetic acid during hydrolysis. In addition to the previous ions, both lipid A fractions resolved signals at *m/z* 1434.0 (loss of 12:0(3-OH) from the main observed structure (*m/z* 1632.1)), *m/z* 1450.0 (loss of 11:0(3-OH) from the main structure), *m/z* 1251.8 (loss of 12:0(3-OH) and 12:0 from the main structure), and *m/z* 1053.6 (loss of 12:0(3-OH)_2_ and 12:0 from the main structure). All these detected ions together with the finding of 12:0(3-OH), 14:0(3-OH), 11:0(3-OH), 12:0 in the composition analysis of KK01 Δ*csaA* support the proposed lipid A structures in Figure 5B.

**Figure 5:**
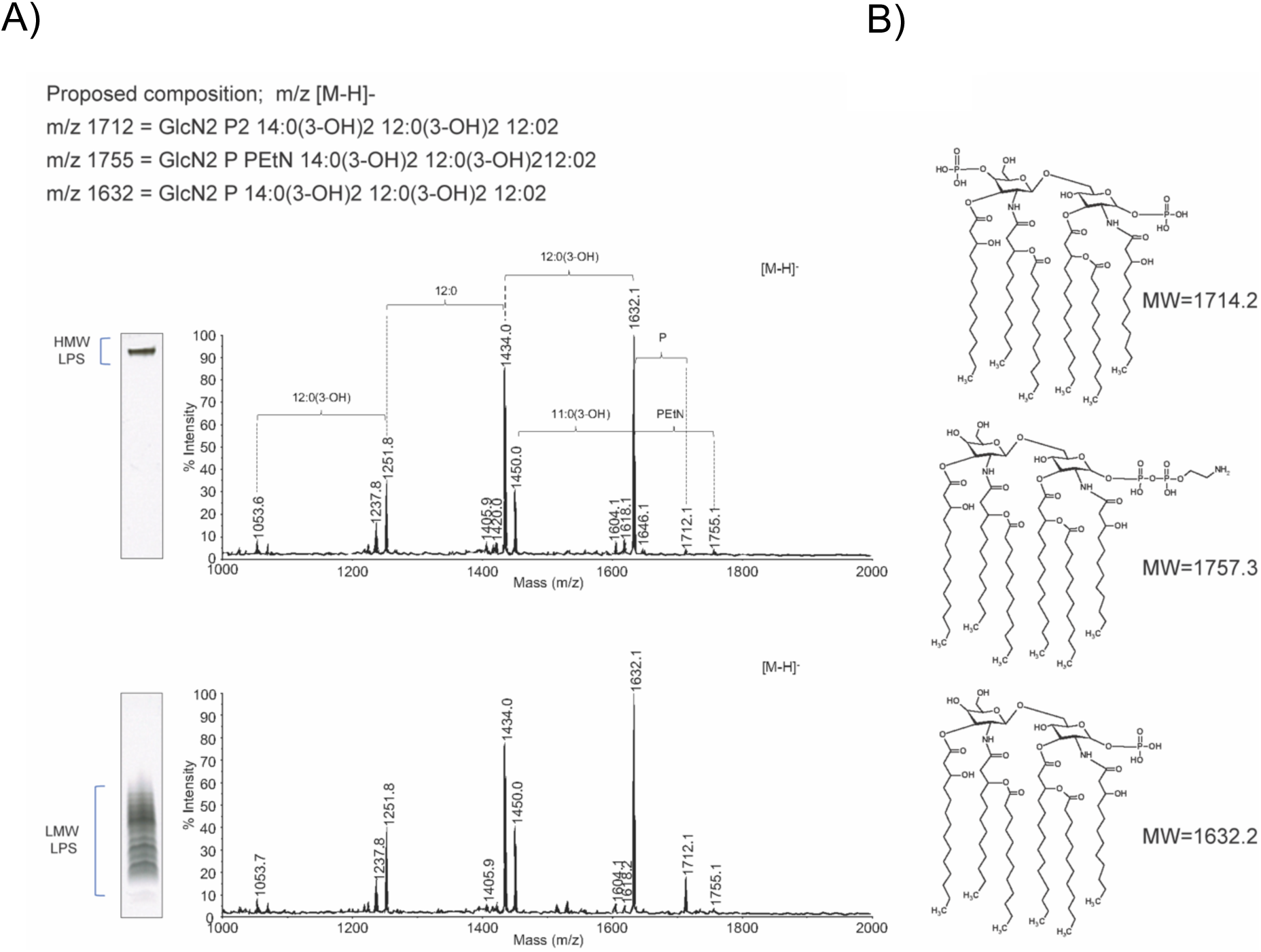
*K. kingae* lipid A can be identified in both HMW and LMW LPS clusters. A) MALDI –TOF MS analysis of the lipid A recovered from the HMW LPS from KK01 Δ*csaA* (Fr 2) (top spectrum), and from LMW LPS of KK01 Δ*csaA* (Fr 3) (bottom spectrum) and proposed lipid A compositions. The spectra were acquired in the negative reflectron ionization mode. The DOC-PAGE insets show the HMW (Fr2) and LMW (Fr 3) LPS. B) Proposed structures of lipid A corresponding to the selected [M-H]^-^ ions observed in MALDI-MS (A).

To better characterize the general structure of the *K. kingae* strain KK01 lipid A from unfractionated LPS, we used a milder lipid A hydrolytic condition on whole bacteria and carried out tandem MS (MS^2^) to confirm the structure of this lipid A molecule. Supplemental Figure 5A shows the mass spectrum obtained from a bacterial solution of KK01 Δ*csaA* after FLAT processing (MS^1^).^26^ Using the precursor ion at *m/z* 1712.12 and 1835.13, the *m/z* difference of 123.01 indicates that the lipid A has a PEtN modification (Supplemental Figure 5A). Tandem MS (MS^2^) was used to confirm the structure. We noted key bond cleavages and fragment ions generated by collision induced dissociation (CID) and FLAT^n^ processing and indicated positions corresponding to each cleavage are in the accompanying structures (Supplemental Figures 5C and 5D). For example, the fragment ion generated by the loss of H_3_PO_4_ from *K. kingae* lipid A corresponds to an ion at *m/z* 1614.14 (theoretical value) and is labeled B_2_ according to accepted nomenclature.^27, 28^ As expected, many other neutral losses were observed, including acyl chains from ester bonds and cross-ring cleavages. Multiple losses of acyl chains from ester bonds, such as *m/z* 1397.97 (B_2_ with 3’α), 1199.81 (B_2_ with 3’α and 3’β), and 999.63 (B_2_ with 3’α, 3 β, and 2’ ε) are shown in Supplemental Figure 5C. The cross-ring cleavage produces ions at *m/z* 690.40 corresponding to ^0,4^A_2_ plus 3’α. These selective dissociations of acyl chains from ester bonds and cross-ring cleavage of the backbone were used to deduce the location of additional acylation groups and modifications of acyl chains. Furthermore, the modification site of lipid A by PEtN was characterized using FLAT^n^. The characteristic ion at *m/z* 987.57 represents the 1-phosphate group modification with PEtN, generated from Y_1_.

Gas chromatography-Flame ionization detection (GC-FOD) analysis of fatty acid content revealed four fatty acids attributed to the lipid A structure. The 12:0(3-OH) and 14:0(3-OH) represent the fatty acids that are ester and amide linked to the glucosamine backbone, respectively. The 12:0 represents the fatty acids that are ester linked to the hydroxyl group at the 3-position. There is a small population of 14:0 fatty acids that are likely in place of 12:0 fatty acids, a common occurrence seen across several lipid A structures (Supplemental Figure 5B). These parallel characterization approaches confirmed the structure of the *K. kingae* lipid A and provided strong evidence that galactan is anchored to LMW LPS.

### The *pamDE* genes are necessary to promote bacterial resistance to polymyxin B- mediated killing

To evaluate the independent functional roles of LPS and galactan, we performed bactericidal assays with polymyxin B at a range of concentrations. Polymyxin B is a cationic antimicrobial peptide that works by displacing the LPS-coordinated metal ions. Previous work by Muñoz *et al.* attributed polymyxin B resistance in *K. kingae* to galactan prior to our knowledge that surface expression of galactan may be dependent on the presence of LMW LPS.^8^ To understand the relative contributions of LPS and galactan to polymyxin B sensitivity, we performed survival assays with strains KK01 Δ*csaA*, KK01 Δ*csaApamABCDE,* KK01 Δ*csaApamABC,* and KK01 Δ*csaApamDE*. As shown in Figure 6, Survival of strains KK01 Δ*csaA* and KK01 Δ*csaApamABC* was not significantly affected by polymyxin B. However, survival of strains KK01 Δ*csaApamABCDE* and KK01 Δ*csaApamDE* was markedly decreased in the presence of polymyxin B (Figure 6). These results establish that the *pamDE* genes are critical for survival in polymyxin B, indicating the key role of LMW LPS and the negligible role of galactan.

**Figure 6:**
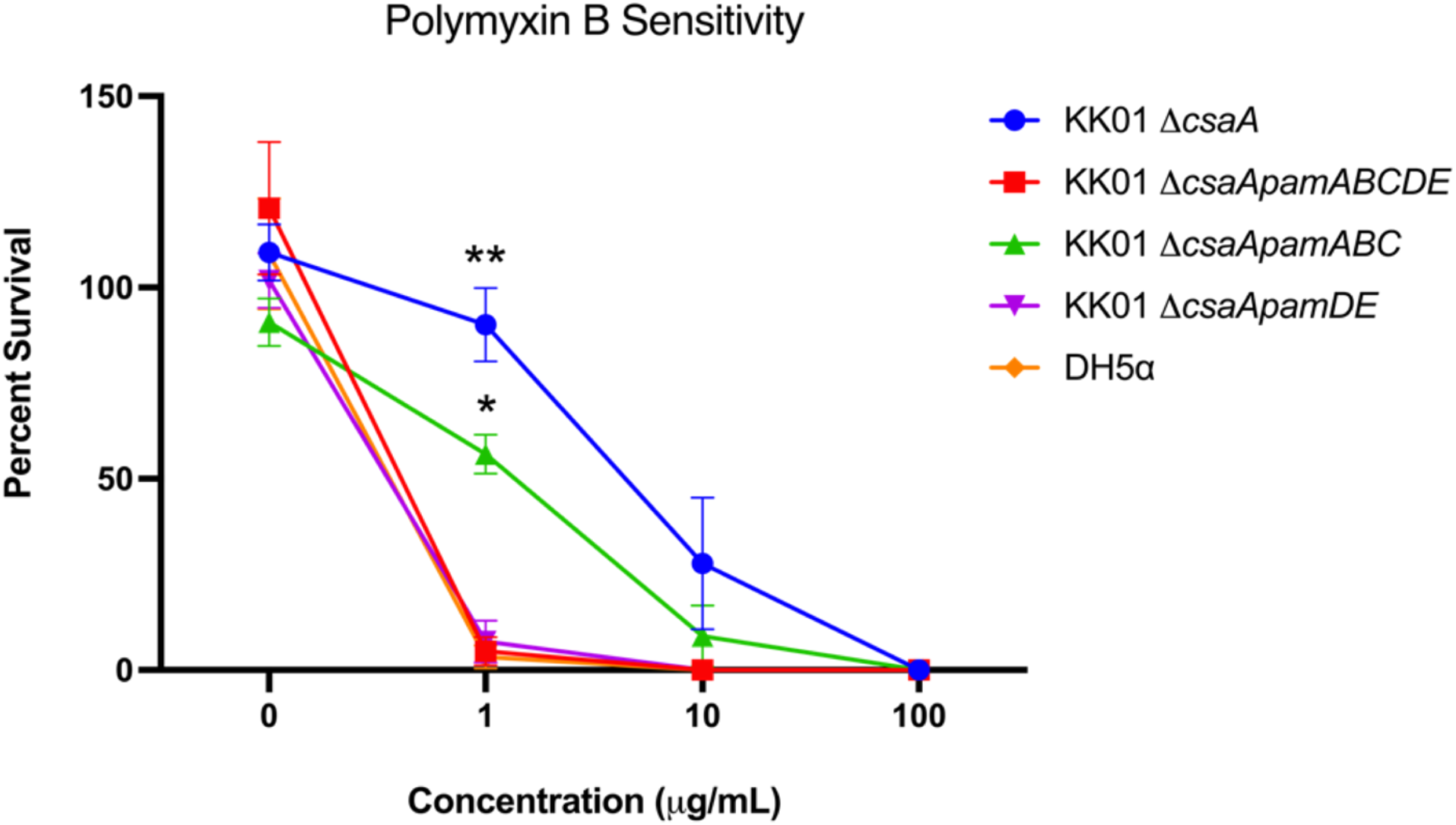
KK01 Δ*csaApamDE* mutants are susceptible to killing by polymyxin B. *K. kingae* strains KK01 Δ*csaA,* Δ*csaApamABCDE,* Δ*csaApamABC,* and Δ*csaApamDE* were challenged with various concentrations of polymyxin B for 30 minutes. CFU were enumerated and survival was determined as a percentage of the inoculum. Data are represented as mean ± SEM from three independent experiments. Statistical significance was determined relative to strain KK01 Δ*csaApamDE* by 2-way ANOVA with Tukey’s correction for multiple comparisons. *, *P < 0.05.* **, *P <* 0.01.

## Discussion

*Kingella kingae* is the leading cause of osteoarticular infections in young children and expresses a collection of surface polysaccharides that contribute to the pathogenesis of disease. The galactan promotes both resistance to serum complement and evasion of neutrophil killing.^8^ Galactan is encoded by the *pamABC* genes that are present in the five-gene *pamABCDE* cluster. In this study we showed that the *pamD* and *pamE* genes in this cluster are required for production of an atypical O-antigen, suggesting that they encode LPS glycosyltransferases. Further analysis by DOC-PAGE and NMR revealed that wild type *K. kingae* produces multiple glycoforms of LPS, including low molecular weight species (LMW LPS) composed primarily of lipid A, core oligosaccharide, and O-antigen and high molecular weight species (HMW LPS) composed of lipid A, core oligosaccharide, O-antigen, and galactan (Figure 7).

**Figure 7:**
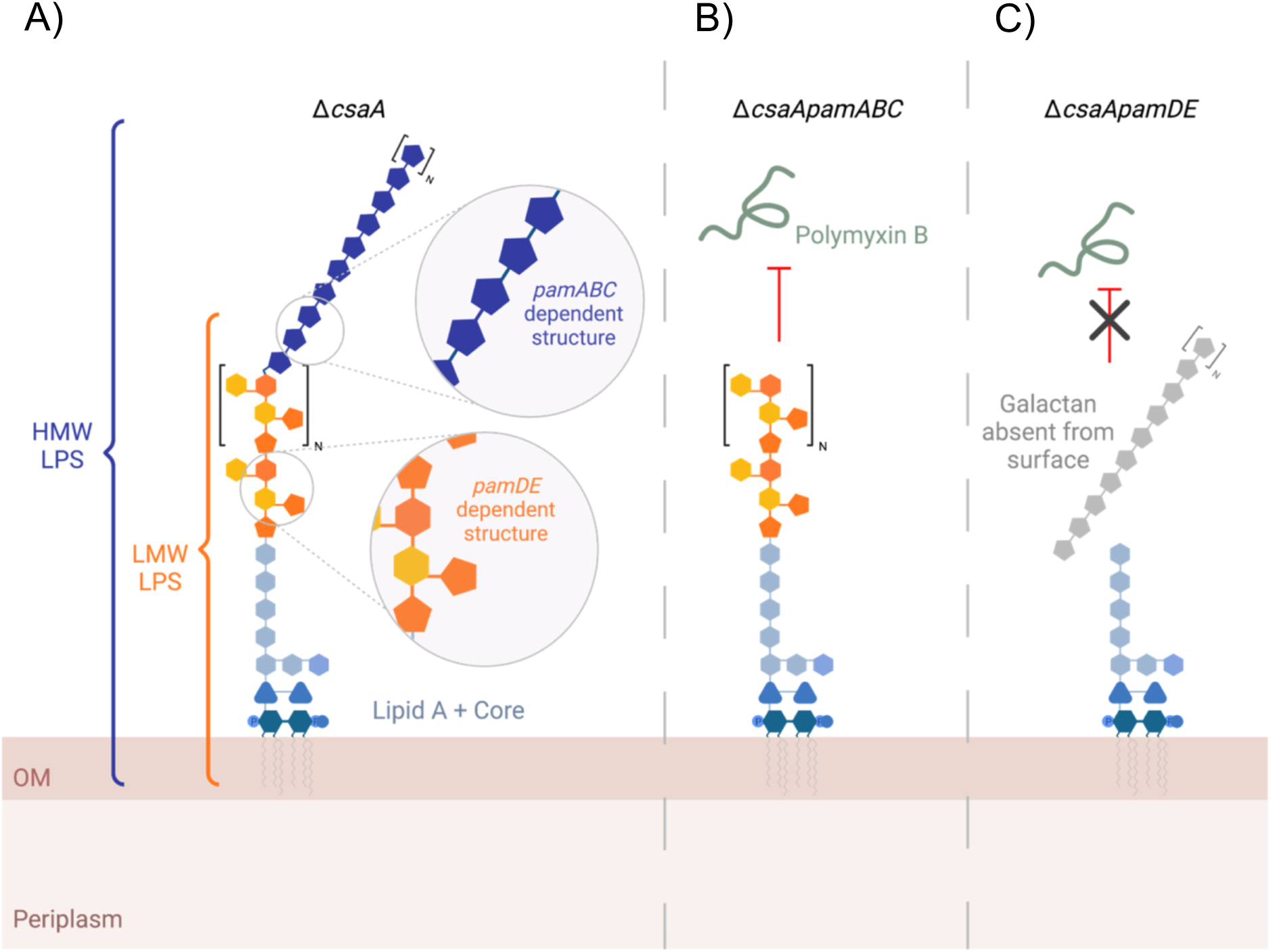
Summary of the relationship between the *K. kingae* galactan and lipopolysaccharide. Schematic representation of the multiple LPS glycoforms expressed by strains KK01 Δ*csaA* (panel A), KK01Δ *csaApamABC* (panel B) and KK01 Δ*csaApamDE* (panel C). LMW LPS glycoforms are composed of lipid A, core oligosaccharide and the *pamDE* dependent O-antigen. Note, the orange shaded O-antigen is depicted as a general repeating unit as the structure is currently unknown. HMW LPS glycoforms are composed of lipid A, core oligosaccharide, and the *pamABC* dependent galactan. Expression of the *pamDE* dependent structure is required for inhibition of polymyxin B mediated killing.

Additional analysis demonstrated that presence of galactan on the bacterial surface is dependent on anchoring to the atypical O-antigen, which in turn is the primary determinant of *K. kingae* resistance to polymyxin B.

NMR and compositional analyses on fractionated LPS and O-PS released from lipid A established that HMW LPS contains a large portion of galactofuranose, suggesting the presence of a novel covalent linkage between the *K. kingae* LPS and galactan. In support of this possibility, we showed that galactan can be detected with a galactan-specific antiserum (GP-19) in an LPS preparation from bacteria with an intact *pam* locus. The galactan used for generation of the galactan-specific antiserum was isolated from the bacterial surface and purified by size exclusion chromatography, making it unlikely that the co-purifying LPS was a contaminant. Fractionation and structural analyses of the HMW LPS and LMW LPS fractions allowed us to identify *K. kingae* lipid A signatures in both fractions, further supporting that the co-purification of LPS and galactan is the result of anchoring of galactan to the atypical O-antigen in LMW LPS. It is important to note that a direct covalent linkage between galactan and LMW LPS could not be observed by NMR because of the complexity of the LMW LPS fraction and possible lability of the Gal*f* linkage during release of the O-PS by acid hydrolysis. However, the biological data and the preliminary NMR analysis presented in this work provide compelling evidence that a covalent linkage exists, uncovering a novel physical relationship between a bacterial exopolysaccharide and LPS.

In other organisms modifications to the O-antigen contribute to the heterogeneity of LPS structures and are recognized as an important driver of host immune recognition.^29–31^ Modification of the O-antigen with acetyl or fucosyl groups is characterized by the transfer of single moieties to individual repeating units and typically occurs in the periplasm by transferase genes present in the major O-antigen gene cluster.^32–34^ Currently there are no other examples of a bacterial LPS with an O-antigen-like structure that is further modified with a very large homopolymeric sugar component. While our data suggest that the transferase required for attachment may be encoded by the *pam* locus, more studies are needed to determine which genes are immediately responsible for creating a covalent linkage between galactan and LMW LPS and to identify the point of attachment.

There are few examples of an association between any component of LPS and an exopolysaccharide. The closest parallel to our observation with *K. kingae* is the expression of M-antigen in *E. coli* K-12. M-antigen is an LPS glycoform in which colanic acid (CA) exopolysaccharide repeat units are polymerized onto the growing LPS molecule with a ligation point at the O-7 of the *L-*glycero-*D*-mannose-heptose in the LPS outer core region.^14^ This is the same residue to which the O-antigen is attached; thus, under CA inducing conditions, a portion of the O-antigen is replaced with CA. Conversely, our data indicate that expression of *pamDE* and a complete LMW LPS is required for surface presence of galactan, suggesting that in *K. kingae* the O-antigen and galactan exopolysaccharide are components of the same LPS molecule.

The LPS genetic machinery is often a good indicator of the mechanism of O-antigen chain length regulation. It is likely that more rigorous genetic mining will be required to identify chain length regulation genes in *K. kingae.* However, the LPS fractionation presented in this work revealed the presence of two isolated modal clusters of LPS species, suggesting that both the HMW LPS and LMW LPS have a narrow distribution of chain lengths.^35^ O-antigen chain length regulation largely occurs by one of two pathways: the ABC-transporter dependent pathway or the *wzx/wzy* pathway.^36^ In *E. coli* and *S. enterica,* the ABC-transporter dependent pathway has been associated with homopolymeric O-antigens.^37, 38^ Similarly, in *P. aeruginosa*, which expresses multiple glycoforms of LPS simultaneously, the ABC-transporter dependent pathway is required for export and regulation of the homopolymeric common polysaccharide antigen (CPA).^39–41^ This pathway begins with initial synthesis of a sugar residue on a lipid adapter on the cytoplasmic leaf of the inner membrane. Once a complete polymer is assembled, it is flipped into the periplasm by the activity of the *wzm/wzt* ABC-transporter.^36, 37, 42^ Once in the periplasm, the O-antigen homopolymer is ligated to the lipid A-core oligosaccharide by the integral membrane protein, WaaL.^37^ Recently, we have identified a putative *waaL* homolog in *K. kingae,* and future analyses will determine if expression of this gene is required for production of HMW LPS or LMW LPS.

Given our current understanding of the homopolymeric nature of galactan and the lack of obvious *wzx/wzy* homologs in the *K. kingae* genome, we hypothesize that HMW LPS may be regulated in an ABC-transporter dependent manner. Preliminary data suggest that the galactan biosynthetic machinery is localized to the cytoplasm, which may support an ABC-transporter dependent mode of LPS assembly. As an additional consideration, it is possible that the *K. kingae* LPS requires two independent mechanisms of chain regulation for the HMW LPS and LMW LPS glycoforms. The chain length of LMW LPS may be regulated in a substrate-mediated manner. In other words, the addition of a critical residue may trigger the addition of the complete galactan polymer, constraining the distribution of chain lengths of LMW LPS. More mutational analyses are required to determine the substrate preference of the *pam* glycosyltransferases. Coupling of this information with a detailed compositional and structural analysis of LMW LPS may provide insight into the mechanism of chain length regulation of the *K. kingae* LPS.

Much work has been done to characterize the function of galactan. Initial studies performed by Bendaoud *et al.* showed that galactan exhibited widespread anti-biofilm properties against biofilms of *Kingella* and other genetically diverse bacteria.^10^ More recently our group examined the role of galactan in resistance to immune mechanisms. These studies demonstrated that galactan is sufficient to prevent complement-mediated killing in the absence of the polysaccharide capsule,^5^ contrasting with observations in other encapsulated organisms, where the capsule plays a dominant role in serum resistance and exopolysaccharides have rarely been implicated.^43–45^ In addition to the unique role of galactan in serum resistance, we found that galactan inhibits neutrophil phagocytosis.^8^ We also reported that galactan mediates resistance to killing by cationic antimicrobial peptides (CAMPs).^8^ It is noteworthy that all previous characterization of galactan has been performed in the context of a full *pamABCDE* locus deletion. In this study, we showed that the deletion of *pamDE* results in the loss of both LMW LPS and surface expression of galactan, a confounding factor that was not considered in earlier work. Our results in this study established that Δ*csaApamDE* mutants are susceptible to polymyxin B killing, while Δ*csaApamABC* mutants remain resistant. CAMPs such as polymyxins function through electrostatic interactions with the gram-negative bacterial outer membrane, which is made negatively charged by the phosphate backbone of the LPS.^46, 47^ Substitution of lipid A with positively charged moieties like PEtN and 4-amino-4-deoxy-*L*-arabinose (*L-*Ara4N) decreases the net negative charge of the membrane and weakens interactions between polymyxins and the membrane, inhibiting bacterial killing.^48, 49^

As an interesting comparison, in *Burkholderia cenocepacia,* a complete inner core oligosaccharide is required for resistance to polymyxins.^49^ In addition to providing the foundation for modifications that increase the net positive charge of the membrane, the inner core oligosaccharide is integral for membrane integrity, which is a prerequisite for antimicrobial resistance.^50^ Here we have shown that the *K. kingae* lipid A is modified with PEtN, which may contribute to polymyxin B resistance, as has been described in the literature.^48, 51, 52^ Additionally, our data suggest that expression of *pamDE* and thus a complete LMW LPS is required for maximal polymyxin B resistance. More work is needed to determine if the mechanism of resistance is through modification of the LPS, steric occlusion of the lipid A backbone by the O-antigen, or crucial membrane integrity that is maintained by LMW LPS. Overall, our data suggest that resistance to CAMPs is mediated by LMW LPS, with little influence of galactan. These results have begun to challenge our understanding of the function of both LPS and galactan, specifically the processes in which these polysaccharides function jointly versus independently.

While this work was performed with our prototype strain KK01 expressing a ➔5)-β-Gal*f*-(1➔ exopolysaccharide, a second exopolysaccharide with the structure ➔3)-β-Gal*f*-(1➔5)-β-Gal*f*-(1➔ has been identified in other clinical isolates of *K. kingae.*^5, 9, 10^ In clinical strain PYKK181, the ➔3)-β-Gal*f*-(1➔5)-β-Gal*f*-(1➔galactan has been shown to promote biofilm dispersal.^10^ However, this exopolysaccharide has not yet been characterized with regard to resistance to serum and neutrophils. Similarly, the diversity of LPS across *K. kingae* isolates is unknown.

Earlier work characterized the *K. kingae* galactan and identified the biosynthetic machinery encoded by the *pam* locus.^7, 9^ In this work we showed that the *pamD* and *pamE* genes are LPS glycosyltransferases required for expression of *K. kingae* LMW LPS and for surface presence of galactan. LPS fractionation and NMR revealed that *K. kingae* HMW LPS contains lipid A, core oligosaccharide, an O-antigen, and galactan, suggesting that galactan is anchored to LMW LPS and revealing novel functions for the *pam* genes. This is the first description of an interaction between *K. kingae* LPS and galactan, highlighting the unique structure of these surface polysaccharides and providing insight into their functions in the pathogenesis of *K. kingae* disease.

## Acknowledgements

This work was supported by the National Institute of Allergy and Infectious Diseases under award 1R01AI121015 (to J.W.S.), and the National Science Foundation Graduate Research Fellowship under awards DGE-1845298 (to N.R.M) and DGE-1321851 (to V.L.M.). Work at the Complex Carbohydrate Research Center was supported by NIH grant R24GM137782-01 to P.A. Work at the University of Maryland-Baltimore was supported by the National Institutes of Health under award R01AI147314 (to R.K.E).

## Author Contributions

Conceptualization, J.W.S., N.R.M, and E.A.P.; Funding acquisition, N.R.M., J.W.S., P.A. and R.K.E.; Formal analysis – N.R.M., E.A.P., A.M., J.V., H.Y., M.S.; Methodology – N.R.M., E.A.P., J.W.S., A.M., J.V., P.A., H.Y., M.S., C.E.C., and R.K.E. Investigation – N.R.M., E.A.P., V.L.M., A.M., J.V., H.Y., M.S., C.E.C.; Project administration – J.W.S., P.A., R.K.E.; Resources – J.W.S., P.A., R.K.E.; Supervision – J.W.S., P.A. and R.K.E.; Validation – N.R.M., E.A.P., J.W.S., V.L.M., A.M., J.V., P.A., H.Y., M.S., C.E.C., and R.K.E.; Visualization – N.R.M., E.A.P., A.M., J.V., H.Y., M.S.; Writing – original draft, N.R.M., E.A.P., J.W.S. A.M., J.V., H.Y., M.S.; Writing – review & editing, N.R.M., E.A.P., J.W.S., V.L.M., A.M., J.V., P.A., H.Y., M.S., C.E.C., and R.K.E.

## Methods

### Bacterial strains and growth conditions

The strains used in this study are listed in Table S3. *K. kingae* and *H. influenzae* strains were stored at –80°C in brain heart infusion (BHI) broth with 20% glycerol. *Escherichia coli* strains were stored at –80°C in Luria-Bertani (LB) broth with 15% glycerol. *K. kingae* strains were grown at 37°C with 5% CO_2_ on chocolate agar. *E. coli* strains were grown at 37°C on LB agar or shaking at 250 rpm in LB broth supplemented with 100 μg/mL ampicillin or 50 μg/mL kanamycin, as appropriate.

### *K. kingae* mutant strain construction

Briefly, plasmid-based gene deletion constructs were created in *E. coli,* linearized, and introduced into *K. kingae* using natural transformation.^53^ Transformants were recovered by selectively plating on chocolate agar plates containing 50 μg/mL kanamycin or 1 μg/mL erythromycin, as appropriate. The introduced mutations were confirmed by PCR and Sanger sequencing. The detailed methods for generating the *K. kingae* mutants used in this study are described in the Supplementary Information Text.

### Surface wash extracts

Bacteria were cultured for 16-20 hours and were suspended in 5 mL PBS to an OD_600_ of 0.8. After 30 minutes of gentle agitation at ambient temperature, the bacteria were removed by centrifugation (30 min at 3220 x *g*), and the supernatant was filtered through a 0.22 μm filter. The resulting supernatant was concentrated to approximately 250 μL over a 50,000 molecular weight cut off (MWCO) Amicon Ultra Centrifugal Filter (MilliporeSigma, Burlington, MA). The samples were supplemented with 2.5 mM MgCl_2_ and 0.1 mM CaCl_2_, treated with 1 unit of DNase I and 10 μg of RNase A for 6 hours at 37°C, followed by treatment with 20 μg of Proteinase K for 16 hrs at 50°C, prior to analysis by silver staining and Western blotting.

### LPS Isolation for DOC-PAGE and ELISA

To isolate LPS, the bacterial strains were grown on chocolate agar for 16-20 hours and resuspended in BHI for the preparation of bacterial lawns. Lawns were grown 16-20 hours on chocolate agar, and the lawns from five plates were pooled by resuspending in PBS. After centrifugation, the pellets were then resuspended in equal amounts of 90% phenol and endotoxin free water, incubated rotating at 70°C for 1 hour, and then centrifuged at 10,000 rpm for 10 minutes to separate the phenolic and aqueous phases. The aqueous phase containing LPS was saved. The phenol layer was reextracted with 500 μL endotoxin free water two more times, and the aqueous layers were pooled. The remaining residual phenol was removed by two 2 mL washes with diethyl ether. The aqueous samples were lyophilized overnight. The lyophilized LPS pellets were resuspended in 1 mL of endotoxin-free water and stored at -20°C. To eliminate contaminating nucleic acids and proteins, LPS samples were treated with 2 units DNase I and 100 μg/mL RNase A for six hours at 37°C and 100 ug/mL proteinase K at 45°C for 16-20 hours.

For ELISA and Western blot analyses, LPS amounts were standardized by Kdo content as described previously.^54^ Briefly, lyophilized LPS samples were resuspended in endotoxin free water. To a 50 μL sample, an equal volume of 0.5N sulfuric acid was added, and the samples were heated at 100°C for 20 minutes. The samples were treated with 36 mM periodate, incubated at room temperature for 10 minutes, and then treated with 0.2 M sodium arsenite and 29 mM thiobarbituric acid. The samples were heated for 8 minutes, cooled to room temperature, and then treated with 1.5 mL of butanol reagent (95% butanol, 5% concentrated HCl). Following centrifugation to separate the phases, the upper butanol layer was removed to a cuvette and promptly read on a spectrophotometer at wavelengths 509 nm and 552 nm. The difference between the two readings was calculated and compared to a Kdo standard curve.

### LPS Isolation for size exclusion chromatography and NMR

The harvested bacterial cells were resuspended in PBS, and the cell suspension was centrifuged for 20 min at 5000 x g at 4°C. The pellets were washed four times until deprived of a viscous supernatant. The crude LPS was obtained from the water phase by applying the hot phenol-water extraction method described by Westphal and Ian.^55^ The water phase was dialyzed against dH_2_O (14,000 MWCO dialysis membrane), then freeze-dried. Nucleic acids and proteins were removed by 12 h treatment with RNase A and DNase I at 37°C, followed by 12 h incubation with proteinase K at 45°C and dialysis (14,000 MWCO, 4°C) against several exchanges of H_2_O. Finally, the samples were ultracentrifuged at 100,000 x *g* at 4°C for 18 h. The LPS pellets were resuspended in water, freeze-dried, and extracted with 9:1 (v/v) ethanol in water at 4°C to remove traces of phospholipids.

### Composition analysis of LPS

The glycosyl composition of LPS was determined by preparation of trimethylsilyl (TMS) methyl glycosides after 18 hours of methanolysis of 300 μg of LPS with 1 M HCl-MeOH at 80°C; in the presence of internal standard of myo-inositol (20 μg).^56^ The TMS method also identified straight-chain and hydroxylated fatty acids constituting LPS (fatty acid methyl esters (FAME) and TMS-FAME, respectively). The TMS and FAME derivatives were analyzed on a Hewlett-Packard HP5890 gas chromatograph, equipped with mass selective detector 5970 MSD using an EC-1 fused silica capillary column (30 m, 0.25 mm I.D.). The oven temperature was 80°C for 2 min, then ramp to 160°C at 20°C/min, and to 200°C at 2°C/min, followed by an increase to 250°C at 10°C/min with an 11-minute hold.

### DOC-PAGE and silver stain analysis

The LPS was analyzed by PAGE using acrylamide gels with deoxycholic acid (DOC) in the running buffer. The DOC-PAGE gels were fixed overnight in 40% ethanol and 5% acetic acid and stained with silver after oxidation with sodium periodate^57^ or with alcian blue followed with classical silver staining as previously described.^58, 59^ The combined alcian blue-silver staining method was used to track and evaluate the LPS elution in the dissociative condition size exclusion chromatography in the presence of DOC.

### Generation of GP-19, α-galactan antiserum

The galactan exopolysaccharide and a recombinant mutant diphtheria toxin carrier protein were purified and conjugated as described in the Supplementary Information Text. The resulting glycoconjugate was sent to Cocalico Biologicals (Stevens, PA) for injection into guinea pig GP-19 using the Ribi adjuvant according to their standard polyclonal antibody production protocol (Cocalico Biologicals IACUC approved project No. 2018-0984), generating the GP-19 antiserum.

### Western blotting

Surface wash extracts, LPS preparations, or digested whole cell sonicates were separated according to the DOC-PAGE method described above and were subsequently transferred to nitrocellulose. Following blocking in 5% skim milk in PBS (blocking buffer), the blots were incubated with antiserum GP-19 diluted 1:1000 in blocking buffer with gentle agitation at ambient temperature for 1 hour or at 4°C for 16-18 hours. Following washing with Tris-buffered saline with 0.1% Tween-20 (TBST), the blots were incubated with a 1:5000 dilution of α-guinea pig-horseradish peroxidase (αGP-HRP) in blocking buffer. After washing with TBST, the blots were exposed to a chemiluminescent HRP substrate, and the blot images were captured using a GBox Chemi:XT4 system (Syngene, Frederick, MD).

### Enzyme-linked immunosorbent assays (ELISAs)

For whole *K. kingae* ELISAs, the strains were cultured 16-20 hours, the bacterial growth was swabbed into PBS to an OD_600_ of 0.1, and 100 μl was added to wells of a 96-well plain polystyrene plate. For *E. coli* whole cell sonicate ELISAs, the digested whole cell sonicates were diluted 1:20 in PBS, and 100 μl was added to the wells of a 96-well MICROLON 200 medium binding plate (Greiner Bio-One, Kremsmünster, Austria). For LPS samples, equivalent quantities were added to 96-well CovaLink (Nunc) plates (ThermoFisher Scientific, Waltham, MA) in 100 μl carbonate buffer (100 mM carbonate, pH 9.6). For all sample types, the plates were processed using the same protocol. Coating was carried out at 4°C for 16-20 hours, and the plates were then washed with TBST and blocked overnight in 2% milk in PBS. GP-19 antiserum dilutions in 2% milk in PBS served as the source of primary antibody for 1 hour at 37°C. The plates were again washed with TBST and were incubated with αGP-HRP (1:2000) for 1 hour at 37°C. The plates were developed with TMB (3, 3’, 5, 5’ - Tetramethylbenzidine) ELISA Peroxidase Substrate (Rockland Immunochemical, Limerick, PA), and absorbance was measured on a plate reader at 655 nm.

### Fractionation of LPS and release of O-PS

HMW LPS was separated from LMW LPS by size exclusion chromatography using a Superdex 75 10/300GL (Cytiva) in dissociative conditions in the presence of 0.25 % sodium deoxycholate, pH 9.2.^60^ The eluting fractions were monitored with refractive index (Shimadzu RID-10A) and with DOC-PAGE stained with alcian blue and silver (see DOC-PAGE method). The O-PS (polysaccharide portion of LPS, including core, O-antigen, and galactan) was released from the lipid A by using 1% acetic acid for 1.5-2h 100°C until the formation of insoluble lipid A precipitate. The lipid A precipitate was removed from the O-PS soluble fraction by initial centrifugation for 25 min at 3500 x *g*. The pellet was washed by adding water, re-suspension, and centrifugation at 100,000 x *g* for 4 h, at 4°C. The soluble O-PS fraction was also ultracentrifuged at 100,000 x *g* for 4 h, 4°C, and the supernatant was freeze-dried. The dry O-PS was dissolved in water and passed through a nylon filter 0.22 µm nylon filter to remove any trace of free lipids, LPS, or lipid A and was used for structural work.

### NMR analysis

Isolated LPS or released O-PS samples were exchanged to D_2_O by lyophilization, and 50 nmol of DSS-*d*_6_ (Cambridge Isotope Laboratories) was added to each sample for chemical shift referencing. NMR data were collected at 25°C on a Varian VNMRS (^1^H, 599.66 MHz) or Bruker Avance III (^1^H, 600.13 MHz) spectrometer, each equipped with a 5 mm cryoprobe. To determine the structure of the released O-PS, 1D ^1^H and 2D COSY, TOCSY, NOESY, HSQC and HMBC spectra were collected. The TOCSY and NOESY spectra were collected with presaturation of the residual water signal, and the HSQC spectrum was acquired with signal multiplicity editing. The mixing times were 70 ms (TOCSY) and 150 ms (NOESY). The homonuclear correlations were collected with ^1^H spectral widths of 4808 Hz, 180 increments and 8–16 scans per increment. The HSQC and HMBC spectra were collected with spectral widths (^1^H and ^13^C) of 7184 and 10556 Hz, 128 increments and 32 (HSQC) or 96 (HMBC) scans per increment. ^1^H and ^13^C chemical shifts were referenced to the respective DSS signals at 0.0 ppm. The NMR data were analyzed in MestreNova 14.1.1.

### MALDI -TOF-MS analysis of lipid A from HMW and LMW LPS

The lipid A released from HMW LPS and LMW LPS fractions that were separated on a Superdex 75 column were analyzed by MALDI-TOF-MS using an Applied Biosystems 4800 Proteomics Analyzer. The lipid A samples were dissolved in a chloroform: methanol solution (3:1; vol./vol.) and mixed with 0.5 M 2,4,6-trihydroxyacetophenone matrix in methanol in a 1:1 ratio. A 0.5 µL volume of the final mixture was applied onto a stainless-steel target plate, and the spectra were acquired in the negative reflector ionization mode ([M-H])^-^.

### Polymyxin B bactericidal assays

Bacterial survival in polymyxin B was determined as described previously.^8^ Briefly, *K. kingae* strains were grown on chocolate agar overnight and resuspended in PBS-G (PBS, 0.1% gelatin). The bacterial suspensions were diluted to a concentration of 4.0 x 10^4^ colony forming units (CFU/mL), 100 μl aliquots were mixed with polymyxin B (Alfa Aesar, Ward Hill, MA) at various physiological concentrations, and samples were incubated at 37°C with 5% CO_2_ for 30 minutes. The reaction was stopped by adding 9 mM MgCl_2_ prior to plating. Serial dilutions of the reactions were plated on chocolate agar plates, and the surviving CFUs were enumerated following overnight incubation. Percent survival was calculated by dividing the recovered CFU counts by the inoculum CFU counts.

### Statistical analysis

Statistical analyses were performed with GraphPad Prism (version 9.2.1) software for Mac (GraphPad Software, San Diego, CA). A *P* value of <0.05 was considered statistically significant. The specific statistical tests used for each experiment are specified in the relevant figure legend.

## References

1. Yagupsky P. Outbreaks of Kingella kingae infections in daycare facilities. Emerg Infect Dis. 2014;20(5):746–753. doi:10.3201/eid2005.131633

2. Yagupsky P, Porsch E, St Geme JW. Kingella kingae: An Emerging Pathogen in Young Children. Pediatrics. Published online 2011. doi:10.1542/peds.2010-1867

3. Yagupsky P. Kingella kingae: carriage, transmission, and disease. Clin Microbiol Rev. 2015;28(1):54–79. doi:10.1128/CMR.00028-14

4. Amit U, Dagan R, Yagupsky P. Prevalence of pharyngeal carriage of kingella kingae in young children and risk factors for colonization. Pediatr Infect Dis J. 2013;32(2):191–193. doi:10.1097/INF.0b013e3182755779

5. Muñoz VL, Porsch EA, Geme JW St. Kingella kingae Surface Polysaccharides Promote Resistance to Human Serum and Virulence in a Juvenile Rat Model. Infect Immun. 2018;86(6):e00100–18. doi:10.1128/IAI.00100-18

6. Kehl-Fie TE, St. Geme JW. Identification and characterization of an RTX toxin in the emerging pathogen Kingella kingae. J Bacteriol. 2007;189(2):430–436. doi:10.1128/JB.01319-06

7. Starr KF, Porsch EA, Seed PC, St Geme III JW. Genetic and Molecular Basis of Kingella kingae Encapsulation. Published online 2016. doi:10.1128/IAI.00128-16

8. Muñoz VL, Porsch EA, St Geme JW. Kingella kingae Surface Polysaccharides Promote Resistance to Neutrophil Phagocytosis and Killing. MBio. 2019;10(3):e00631–19. doi:10.1128/mBio.00631-19

9. Starr KF, Porsch EA, Heiss C, Black I, Azadi P, St. Geme JW. Characterization of the Kingella kingae Polysaccharide Capsule and Exopolysaccharide. Adler B, ed. PLoS One. 2013;8(9):e75409. doi:10.1371/journal.pone.0075409

10. Bendaoud M, Vinogradov E, Balashova N V, Kadouri DE, Kachlany SC, Kaplan JB. Broad-Spectrum Biofilm Inhibition by Kingella kingae Exopolysaccharide. J Bacteriol. 2011;193(15):3879–3886. doi:10.1128/JB.00311-11

11. Kessler NG, Caraballo Delgado DM, Shah NK, Dickinson JA, Moorea SD. Exopolysaccharide anchoring creates an extreme resistance to sedimentation. J Bacteriol. 2021;203(11). doi:10.1128/JB.00023-21/ASSET/6E965858-0EE3-42F3-8822-33FC8DE6F275/ASSETS/IMAGES/LARGE/JB.00023-21-F0008.JPG

12. El-Kazzaz W, Morita T, Tagami H, Inada T, Aiba H. Metabolic block at early stages of the glycolytic pathway activates the Rcs phosphorelay system via increased synthesis of dTDP-glucose in Escherichia coli. Mol Microbiol. 2004;51(4):1117–1128. doi:10.1046/j.1365-2958.2003.03888.x

13. Rinno,’ J, Golecki JR, Mayer’c AH. Localization of Enterobacterial Common Antigen: Immunogenic and Nonimmunogenic Enterobacterial Common Antigen-Containing Escherichia coli. J Bacteriol. 1980;141(2):814–821.

14. Meredith TC, Mamat U, Kaczynski Z, Lindner B, Holst O, Woodard RW. Modification of lipopolysaccharide with colanic acid (M-antigen) repeats in Escherichia coli. J Biol Chem. 2007;282(11):7790–7798. doi:10.1074/jbc.M611034200

15. Gozdziewicz TK, Lugowski C, Lukasiewicz J. First evidence for a covalent linkage between enterobacterial common antigen and lipopolysaccharide in Shigella sonnei phase II ECALPS. J Biol Chem. 2014;289(5):2745–2754. doi:10.1074/jbc.M113.512749

16. Bertani B, Ruiz N. Function and Biogenesis of Lipopolysaccharides. Published online 2019. doi:10.1128/ecosalplus.ESP-0001–2018

17. Falchi FA, Maccagni EA, Puccio S, et al. Mutation and suppressor analysis of the essential lipopolysaccharide transport protein LptA reveals strategies to overcome severe outer membrane permeability defects in Escherichia coli. J Bacteriol. 2018;200(2). doi:10.1128/JB.00487-17

18. Muheim C, Götzke H, Eriksson AU, et al. Increasing the permeability of Escherichia coli using MAC13243. doi:10.1038/s41598-017-17772-6

19. Ram S, Gulati S, Lewis LA, et al. A Novel Sialylation Site on Neisseria gonorrhoeae Lipooligosaccharide Links Heptose II Lactose Expression with Pathogenicity. Infect Immun. 2018;86(8):e00285–18. doi:10.1128/IAI.00285-18

20. Grossman N, Schmetz MA, Foulds J, et al. Lipopolysaccharide size and distribution determine serum resistance in Salmonella montevideo. J Bacteriol. 1987;169(2):856–863. doi:10.1128/JB.169.2.856-863.1987

21. Domínguez-Medina CC, Pérez-Toledo M, Schager AE, et al. Outer membrane protein size and LPS O-antigen define protective antibody targeting to the Salmonella surface. Nat Commun 2020 111. 2020;11(1):1-11. doi:10.1038/s41467-020-14655-9

22. Goebel EM, Wolfe DN, Elder K, Stibitz S, Harvill ET. O antigen protects Bordetella parapertussis from complement. Infect Immun. 2008;76(4):1774–1780. doi:10.1128/IAI.01629-07

23. Park BS, Lee J-O. Recognition of lipopolysaccharide pattern by TLR4 complexes. Exp Mol Med. 2013;45(12):e66. doi:10.1038/emm.2013.97

24. Li Y, Powell DA, Shaffer SA, et al. LPS remodeling is an evolved survival strategy for bacteria. Proc Natl Acad Sci U S A. 2012;109(22):8716–8721. doi:10.1073/PNAS.1202908109/-/DCSUPPLEMENTAL

25. Nichols WA, Gibson BW, Melaugh W, Lee NAG, Sunshine M, Apicella MA. Identification of the ADP-L-glycero-D-manno-heptose-6-epimerase (rfaD) and heptosyltransferase II (rfaF) biosynthesis genes from nontypeable Haemophilus influenzae 2019. Infect Immun. 1997;65(4):1377. doi:10.1128/iai.65.4.1377-1386.1997

26. Yang H, Jackson SN, Woods AS, Goodlett DR, Ernst RK, Scott AJ. Streamlined Analysis of Cardiolipins in Prokaryotic and Eukaryotic Samples Using a Norharmane Matrix by MALDI-MSI. J Am Soc Mass Spectrom. 2020;31(12):2495–2502. doi:10.1021/jasms.0c00201

27. Costello CE, Vath JE. Tandem mass spectrometry of glycolipids. Methods Enzymol. 1990;193(C):738–768. doi:10.1016/0076-6879(90)93448-T

28. Jones JW, Cohen IE, Tureek F, Goodlett DR, Ernst RK. Comprehensive Structure Characterization of Lipid A Extracted from Yersinia pestis for Determination of its Phosphorylation Configuration. J Am Soc Mass Spectrom. 2010;21(5):785–799. doi:10.1016/j.jasms.2010.01.008

29. Wang M, Arbatsky NP, Xu L, Shashkov AS, Wang L, Knirel YA. O antigen of franconibacter pulverisG3872 (O1) is a 4-deoxy-D-arabino-hexose-containing polysaccharide synthesized by the ABC-transporter-dependent pathway. Microbiol (United Kingdom). 2016;162(7):1103–1113. doi:10.1099/mic.0.000307

30. Lerouge I, Vanderleyden J. O-antigen structural variation: mechanisms and possible roles in animal/plant–microbe interactions. FEMS Microbiol Rev. 2002;26(1):17–47. doi:10.1111/j.1574-6976.2002.tb00597.x

31. Kintz E, Scarff JM, DiGiandomenico A, Goldberg JB. Lipopolysaccharide O-antigen chain length regulation in Pseudomonas aeruginosa serogroup O11 strain PA103. J Bacteriol. 2008;190(8):2709–2716. doi:10.1128/JB.01646-07

32. Kubler-Kielb J, Vinogradov E, Chu C, Schneerson R. Acetylation of the O-specific polysaccharide isolated from Shigella flexneri serotype 2a. Carbohydr Res. 2007;342(3-4):643–647. https://www.ncbi.nlm.nih.gov/pmc/articles/PMC3624763/pdf/nihms412728.pdf

33. Pearson CR, Tindall SN, Herman R, et al. Acetylation of surface carbohydrates in bacterial pathogens requires coordinated action of a two-domain membrane-bound acyltransferase. MBio. 2020;11(4):1–19. doi:10.1128/mBio.01364-20

34. Naumenko OI, Zheng H, Xiong Y, et al. Studies on the O-polysaccharide of Escherichia albertii O2 characterized by non-stoichiometric O-acetylation and non-stoichiometric side-chain L-fucosylation. Carbohydr Res. 2018;461:80–84. doi:10.1016/j.carres.2018.02.013

35. Whitfield C, Williams DM, Kelly SD. Lipopolysaccharide O-antigens-bacterial glycans made to measure. J Biol Chem. 2020;295(31):10593–10609. doi:10.1074/jbc.REV120.009402

36. Samuel G, Reeves P. Biosynthesis of O-antigens: genes and pathways involved in nucleotide sugar precursor synthesis and O-antigen assembly. doi:10.1016/j.carres.2003.07.009

37. Greenfield LK, Whitfield C. Synthesis of lipopolysaccharide O-antigens by ABC transporter-dependent pathways. Carbohydr Res. 2012;356:12–24. doi:10.1016/j.carres.2012.02.027

38. DebRoy C, Fratamico PM, Yan X, et al. Comparison of O-antigen gene clusters of all O-serogroups of Escherichia coli and proposal for adopting a new nomenclature for O-typing. PLoS One. 2016;11(1):1–13. doi:10.1371/journal.pone.0147434

39. Rocchetta HL, Lam JS. Identification and functional characterization of an ABC transport system involved in polysaccharide export of A-band lipopolysaccharide in Pseudomonas aeruginosa. J Bacteriol. 1997;179(15):4713–4724. doi:10.1128/JB.179.15.4713-4724.1997

40. Abeyrathne PD, Daniels C, Poon KKH, Matewish MJ, Lam JS. Functional characterization of WaaL, a ligase associated with linking O-antigen polysaccharide to the core of Pseudomonas aeruginosa lipopolysaccharide. J Bacteriol. 2005;187(9):3002–3012. doi:10.1128/JB.187.9.3002-3012.2005

41. Wang S, Hao Y, Lam JS, et al. Biosynthesis of the common polysaccharide antigen of Pseudomonas aeruginosa PAO1: Characterization and role of GDP-D-Rhamnose: GlcNAc/GalNAc-diphosphate-lipid α1,3-D-rhamnosyltransferase WbpZ. J Bacteriol. 2015;197(12):2012–2019. doi:10.1128/JB.02590-14

42. Sperandeo P, Martorana AM, Polissi A. Lipopolysaccharide biogenesis and transport at the outer membrane of Gram-negative bacteria. Biochim Biophys Acta - Mol Cell Biol Lipids. 2017;1862(11):1451–1460. doi:10.1016/j.bbalip.2016.10.006

43. Jones CJ, Wozniak DJ, Joanna Goldberg EB. Psl Produced by Mucoid Pseudomonas aeruginosa Contributes to the Establishment of Biofilms and Immune Evasion. doi:10.1128/mBio.00864-17

44. Mishra M, Byrd MS, Sergeant S, et al. Pseudomonas aeruginosa Psl polysaccharide reduces neutrophil phagocytosis and the oxidative response by limiting complement-mediated opsonization. Cell Microbiol. 2012;14(1):95–106. doi:10.1111/j.1462-5822.2011.01704.x

45. Miajlovic H, Cooke NM, Moran GP, Rogers TRF, Smith SG. Response of extraintestinal pathogenic Escherichia coli to human serum reveals a protective role for Rcs-regulated exopolysaccharide colanic acid. Infect Immun. 2014;82(1):298–305. doi:10.1128/IAI.00800-13

46. Lai Y, Gallo RL. AMPed up immunity: how antimicrobial peptides have multiple roles in immune defense. Trends Immunol. 2009;30(3):131–141. doi:10.1016/j.it.2008.12.003

47. Clifton LA, Ciesielski F, Skoda MWA, Paracini N, Holt SA, Lakey JH. The Effect of Lipopolysaccharide Core Oligosaccharide Size on the Electrostatic Binding of Antimicrobial Proteins to Models of the Gram Negative Bacterial Outer Membrane. Published online 2016. doi:10.1021/acs.langmuir.6b00240

48. Tamayo R, Choudhury B, Septer A, Merighi M, Carlson R, Gunn JS. Identification of cptA, a PmrA-regulated locus required for phosphoethanolamine modification of the Salmonella enterica serovar typhimurium lipopolysaccharide core. J Bacteriol. 2005;187(10):3391–3399. doi:10.1128/JB.187.10.3391-3399.2005

49. Loutet SA, Flannagan RS, Kooi C, Sokol PA, Valvano MA. A complete lipopolysaccharide inner core oligosaccharide is required for resistance of Burkholderia cenocepacia to antimicrobial peptides and bacterial survival in vivo. J Bacteriol. 2006;188(6):2073–2080. doi:10.1128/JB.188.6.2073-2080.2006

50. Rahaman SO, Mukherjee J, Chakrabarti A, Pal S. Decreased Membrane Permeability in a Polymyxin B-Resistant Escherichia Coli Mutant Exhibiting Multiple Resistance to L-Lactams as Well as Aminoglycosides. doi:10.1111/j.1574-6968.1998.tb12955.x

51. Wright JC, Hood DW, Randle GA, et al. lpt6, a gene required for addition of phosphoethanolamine to inner-core lipopolysaccharide of Neisseria meningitidis and Haemophilus influenzae. J Bacteriol. 2004;186(20):6970–6982. doi:10.1128/JB.186.20.6970-6982.2004

52. Sorensen M, Chandler CE, Gardner FM, et al. Rapid microbial identification and colistin resistance detection via MALDI-TOF MS using a novel on-target extraction of membrane lipids. Sci Rep. 2020;10(1):1–9. doi:10.1038/s41598-020-78401-3

53. Kehl-Fie TE, St Geme JW. Identification and characterization of an RTX toxin in the emerging pathogen Kingella kingae. J Bacteriol. 2007;189(2):430–436. doi:10.1128/JB.01319-06

54. Hancock RE. (University of BC. KDO Assay. Hancock Laboratory Methods. Published 1999. Accessed October 3, 2022. http://cmdr.ubc.ca/bobh/method/kdo-assay/

55. Westphal O, Jann K. Bacterial lipopolysaccharides. Methods Carbohydr Chem. 1965;5:83–91.

56. York WS, Darvill AG, McNeil M, Stevenson TT, Albersheim P. Isolation and characterization of plant cell walls and cell wall components. Methods Enzymol. 1986;118(C):3–40. doi:10.1016/0076-6879(86)18062-1

57. Krauss JH, Weckesser J, Mayer H. Electrophoretic analysis of lipopolysaccharides of purple nonsulfur bacteria. Int J Syst Bacteriol. 1988;38(2):157–163. doi:10.1099/00207713-38-2-157/CITE/REFWORKS

58. Corzo J, Pérez-Galdona R, León-Barrios M, Gutiérrez-Navarro AM. Alcian blue fixation allows silver staining of the isolated polysaccharide component of bacterial lipopolysaccharides in polyacrylamide gels. Electrophoresis. 1991;12(6):439–441. doi:10.1002/ELPS.1150120611

59. Muszynski A, Laus M, Kijne JW, Carlson RW. Structures of the lipopolysaccharides from Rhizobium leguminosarum RBL5523 and its UDP-glucose dehydrogenase mutant (exo5). Glycobiology. 2011;21(1):55–68. doi:10.1093/GLYCOB/CWQ131

60. Reuhs BL, Kim JS, Badgett A, Carlson RW. Production of cell-associated polysaccharides of Rhizobium fredii USDA205 is modulated by apigenin and host root extract. Mol Plant Microbe Interact. 1994;7(2):240–247. doi:10.1094/MPMI-7-0240

