## Supplemental Information for "Surface anchoring of the *Kingella kingae* galactan is dependent on the lipopolysaccharide O-antigen"

### Supplementary Information

#### Supplementary Materials and Methods

*K. kingae* mutant strain construction

The KK01 Δ*csaA* strain used in this study differs from the previously described mutant[1] in that the *aphA3* cassette was excised using the spot transformation procedure,[2] resulting in an unmarked deletion. Briefly, serial 4-fold dilutions of the *aphA3*-marked KK01Δ*csaA* mutant in BHI supplemented with 50 mM $MgCl_2$ were incubated with a linearized derivative of plasmid pUC19*csaA,*[1] pUC19*csaA*::unmarked, containing regions with 5' (upstream) and 3' (downstream) homology to the *csaA* open reading frame (ORF) but lacking the *aphA3* marker cassette between the two flanking regions. The transformation reactions were plated on chocolate agar, and single colonies were picked and subjected to two rounds of single colony purification, at which point they were also struck on chocolate agar with 50 µg/mL kanamycin. The colonies that lost kanamycin resistance were further screened by PCR and Sanger sequencing to confirm presence of the unmarked *csaA* deletion. Deletion of the capsule synthesis locus had no effect on growth on chocolate agar or on production of galactan.

To create the *pamABC* deletion construct, an approximately 1000-bp fragment immediately upstream of the *pamA* ORF and an approximately 1000-bp fragment immediately downstream of the *pamC* ORF were separately amplified from strain KK01 with primers *pamA5'_F/pamA5'_R,* and *pamC3'_F*/*pamC3'_R*, respectively (see Table S4 for primer sequences). The *pamA5'* fragment was digested with EcoRI/KpnI and ligated into EcoRI/KpnI-digested pUC19, generating pUC19-*pamA5'*. The *pamC3'* fragment was digested with BamHI/HindIII and was ligated into BamHI/HindIII-digested pUC19-*pamAup*, generating pUC19-*pamA5':pamC3'*. The *aphA3* kanamycin resistance cassette was amplified with primers *aphA3_F*/*aphA3*_R from plasmid pFalcon2, digested with KpnI/BamHI, and ligated into KpnI/BamHI-digested pUC-*pamA5':pamC3'*, generating plasmid pΔ*pamABC*. The same approach was used to generate pΔ*pamDE* (using primers *pamD5'_F/pamD5'_R*, *pamE5'_F/pamE5'_R*, and *aphA3_F*/*aphA3*_R) and pΔ*rfaF* (using primers *rfaF*5'_F/*rfaF*5'_R, *rfaF*3'_F/*rfaF*3'_R, and *ermC_F/ermC_R*), with the exception that the *ermC* erythromycin resistance cassette (amplified from pIDN4) was used as the marker for pΔ*rfaF* instead of *aphA3*. The resulting plasmids, along with plasmid pUC19*pam*::*ermC*[3] used to delete the entire *pamABCDE* locus, were linearized and separately transformed into KK01 Δ*csaA* via natural transformation.

<u>Galactan production in *E. coli*</u>

The *pamABC* genes were PCR amplified as a single fragment, starting with the predicted *pamA* start codon, with primers *pamABC*_F and *pamABC*_R, digested with EcoRI/KpnI, and ligated into EcoRI/KpnI-digested pTrc99a, generating plasmid pTrc99a-*pamABC*. The *pamDE* genes were PCR amplified as a single fragment, starting approximately 150 bp upstream of the predicted *pamD* start codon, with primers *pamDE*_F and *pamDE*_R, and were ligated into SalI/BamHI-digested pACYC184 using the NEB Hifi Assembly Kit (New England Biolabs, Ipswich, MA), generating plasmid pACYC184-*pamDE*. Expression of *pamABC* in pTrc99a-*pamABC* is under control of the ITPG-inducible Trc promoter, while expression of *pamDE* in pACYC184-*pamDE* is under control of the predicted native *K. kingae pamD* promoter. Both plasmids were confirmed to be correct by restriction digestion and Sanger sequencing and were then transformed into *E. coli* strain JM109.

To examine galactan production in *E. coli*, strains JM109/pTrc99a, JM109/pTrc99a-*pamABC*, JM109/pACYC184, JM109/pACYC184-*pamDE*, and JM109/pTrc99a-*pamABC*+pACYC-*pamDE* were cultured overnight at 37°C shaking in LB supplemented with 100 µg/mL ampicillin (to select for pTrc99A) and/or 20 µg/mL chloramphenicol (to select for pACYC184) as appropriate. After overnight growth, the cultures were back-diluted 1:100 in LB containing the appropriate antibiotic(s) and were cultured at 37°C shaking until the $OD_{600}$ reached ~0.4, at which point IPTG was added to a final concentration of 0.4 mM. After 3 hours of growth under inducing conditions, the bacteria were collected by centrifugation. The media was removed, and the pellet was resuspended to an $OD_{600}$ of 1.0 in PBS. Three milliliters of the bacterial resuspension was pelleted, resuspended in 0.5 ml PBS, and sonicated 3 x 30 sec. $MgCl_2$ (2.5 mM) and $CaCl_2$ (0.1 mM) were added to the whole cell sonicate, followed by treatment with DNase I (5 units) and RNase A (100 µg) for 6 hours at 37°C. Proteinase K (100 ug) was then added, and the digestion was carried out at 45°C for an additional 12-16 hours. The resulting digested whole cell sonicates were then processed for Western blotting or ELISA analyses.

<u>Generation of GP19, α-galactan antiserum</u>

*Galactan purification.* Strain KK01 ∆*csaA* was grown 16-20 hours on chocolate agar, and the bacterial growth was suspended in BHI broth. Using a sterile swab, 40 chocolate plates were inoculated from the BHI suspension to generate a lawn of growth on each plate. After overnight growth, the 40 lawn plates were swabbed to collect the bacterial mass into 160 ml PBS using sterile polyester tipped swabs, and this suspension was then gently agitated at ambient temperature for 30 min. After centrifugation at 6,700 x *g* for 20 min, the supernatant was filtered through a 0.22 μm filter, dialyzed against 5 L of water over two days with one water change (10 L total dialysis) using 10,000 MWCO dialysis tubing, flash frozen, and lyophilized. The lyophilized material was suspended in 10 ml PBS, extracted twice with Tris-saturated phenol, dialyzed against 5 L of water with 3 water changes over 2 days (20 L total dialysis), flash frozen, and lyophilized. The lyophilized material was resuspended in PBS supplemented with 2.5 mM $MgCl_2$ and 0.1 mM $CaCl_2$, treated with 20 units of DNase I and 200 μg of RNase A for 16 hours at 37°C, and then treated with 500 μg of proteinase K for 16 hours at 45°C. The sample was then filtered through a 0.22 μm filter and separated over a HiLoad 16/600 200 pg size exclusion column (Cytiva, Marlborough, MA). The fractions were analyzed by silver staining for presence of galactan. The fractions containing galactan were pooled, and total polysaccharide was precipitated by addition of 95% ethanol to a final concentration of 75%. The precipitated polysaccharide was collected by centrifugation at 16,000 x *g* for 20 min and washed three times with 75% ethanol. The resulting material was dissolved in water, and the sugar content was quantified using the phenol-sulfuric acid assay for hexoses and pentoses with galactose as the standard.[4] An aliquot was then subjected to glycosyl composition analysis.

*DT 51E/148K purification.* To generate a high-titer antiserum to galactan, we chose to generate a glycoconjugate of galactan and an immunogenic carrier protein to serve as the antigen for antibody production in a guinea pig. We selected a 6x His-tagged inactive mutant diphtheria toxin (DT 51E/148K) as the carrier protein. *E. coli* DH5α containing plasmid pET-22b DT 51E/148K, which was a gift from John Collier (Addgene plasmid # 11081; http://n2t.net/addgene:11081; RRID:Addgene_11081), was cultured overnight at 37°C shaking and back diluted 1:100 in LB ampicillin (100 µg/mL) in 1 L. When the culture reached an $OD_{600}$ of ~0.4, IPTG was added to a final concentration of 0.4 mM, and the culture was moved to 30°C shaking for three hours for protein induction. The bacteria were collected by centrifugation, resuspended in 50 mL His binding buffer (20 mM sodium phosphate pH 7.4, 500 mM NaCl, 40 mM imidazole) containing complete EDTA-free Protease Inhibitor Cocktail (MilliporeSigma), and sonicated 3 x 30 seconds. The sonicate was clarified by centrifugation at 12,000 x *g* for 30 min at 4°C. After filtration through a 0.22 µM filter, the clarified lysate was applied to a 5 mL HisTrap column (Cytiva), washed with His binding buffer, and eluted with His binding buffer containing 500 mM imidazole. Fractions containing DT 51E/148K were pooled, concentrated, and buffer exchanged over a 30,000 MWCO Amicon filter to remove excess imidazole, and further purified over a HiLoad 16/600 200 pg size exclusion column. Fractions containing DT 51E/148K were pooled, concentrated to 2 mg/mL over a 30,000 MWCO Amicon filter, and dialyzed into 0.9% NaCl for conjugation.

*Galactan-DT 51E/148K conjugation.* The galactan was covalently linked to DT 51E/148K using an adapted method of 1-cyano-4-dimethylaminopyridinium tetrafluroborate (CDAP) conjugation.[5] One-hundred microliters of CDAP in acetonitrile (100 mg/mL) was slowly added to 2 mL of 1 mg/mL galactan. After 30 seconds of incubation at ambient temperature, 100 µl of aqueous 0.2 M triethylamine was added. Next, 1 mL of 2 mg/mL DT 51E/148K in 0.25 M HEPES pH 8.2 was added and gently mixed. The conjugation reaction was incubated at ambient temperature for 24 hours, extensively diafiltered with 0.9% NaCl through a 100,000 MWCO Amicon filter to remove the unconjugated carrier protein and conjugation chemicals and was analyzed by SDS-PAGE with silver and Coomassie blue staining.

The resulting glycoconjugate was sent to Cocalico Biologicals (Stevens, PA) for injection into guinea pig GP-19 using the Ribi adjuvant according to their standard polyclonal antibody protocol (Cocalico Biologicals IACUC approved project number 2018-0984), generating the GP-19 antiserum.

FLAT lipid A isolation, MALDI-TOF and MS/MS

To isolate lipid A from unfractionated LPS, bacterial lawns were prepared as described above. Pellets were resuspended in 1 mL of endotoxin free water, and 1 µL was used for analysis. The sample was deposited on the ITO glass slide and allowed to dry. FLAT was conducted as described previously.[6] Briefly, 1 µL of the prepared citrate buffer solution (0.2 M citric acid, 0.1 M trisodium citrate, pH 3.5) was deposited onto the sample spot on the ITO slide. The plate was incubated in a humidified, closed glass chamber for 30 min at 110°C. After heating, the ITO slide was removed from the chamber and cooled, and the plate was thoroughly washed several times with water using a pipettor and left to dry on the laboratory bench. In all cases, 10 mg/mL of norharman (NRM)[6] in 1:2 MeOH:$CHCl_3$ (v:v) was used for lipid A detection. NRM solution (1 µL) was deposited on the sample spot.

A Bruker MALDI (tims TOF) MS was used for $FLAT^n$ experiments and was equipped with a dual ESI/MALDI source with a SmartBeam 3D 10 KHz frequency tripled Nd:YAG laser (355 nm). The system was operated in "qTOF" mode (TIMS deactivated). Ion transfer tuning was used with the following parameters: Funnel 1 RF: 440.0 Vpp, Funnel 2 RF: 490.0 Vpp, Multipole RF 490.0 Vpp, is CID Energy: 0.0 eV, and Deflection Delta: -60.0 V. Quadrupole has been used with the following values for MS mode: Ion Energy: 4.0 eV and Low Mass 700.00 *m/z*. Collision cell activation of ions used the following values for MS mode: Collision Energy: 9.0 eV and Collision RF:3900.0 Vpp. In the MS/MS mode, the precursor ion was chosen by typing targeted *m/z* value including two digits to the right of the decimal point. Typical isolation width and collision energy were set to 4 *m/z* and 100 eV, respectively.

Data processing

All MALDI (timsTOF) MS and MS/MS data were visualized using mMass (Ver 5.5.0).[7] Peak picking was conducted in mMass. Identification of all fragment ions were determined based on Chemdraw Ultra (Ver10.0).

Lipid A fatty acid analysis

LPS fatty acid content was measured via gas chromatography coupled to flame ionization detection (GC-FID) after acid hydrolysis, methylation, and hexane extraction.[8] Briefly, LPS was isolated as described above, and cleavage of the ester and amide bonds linking the acyl chains to the glucosamine backbone was achieved by incubating LPS in the presence of methanolic HCl, subsequently converting fatty acids into fatty acid methyl esters. These fatty acid methyl esters were then extracted using hexane and analyzed via GC-FID. Peak assignments were made based on the retention time of FAME standards. Quantitation of FAME peaks was performed using a pentadecanoic acid (C15) internal standard.

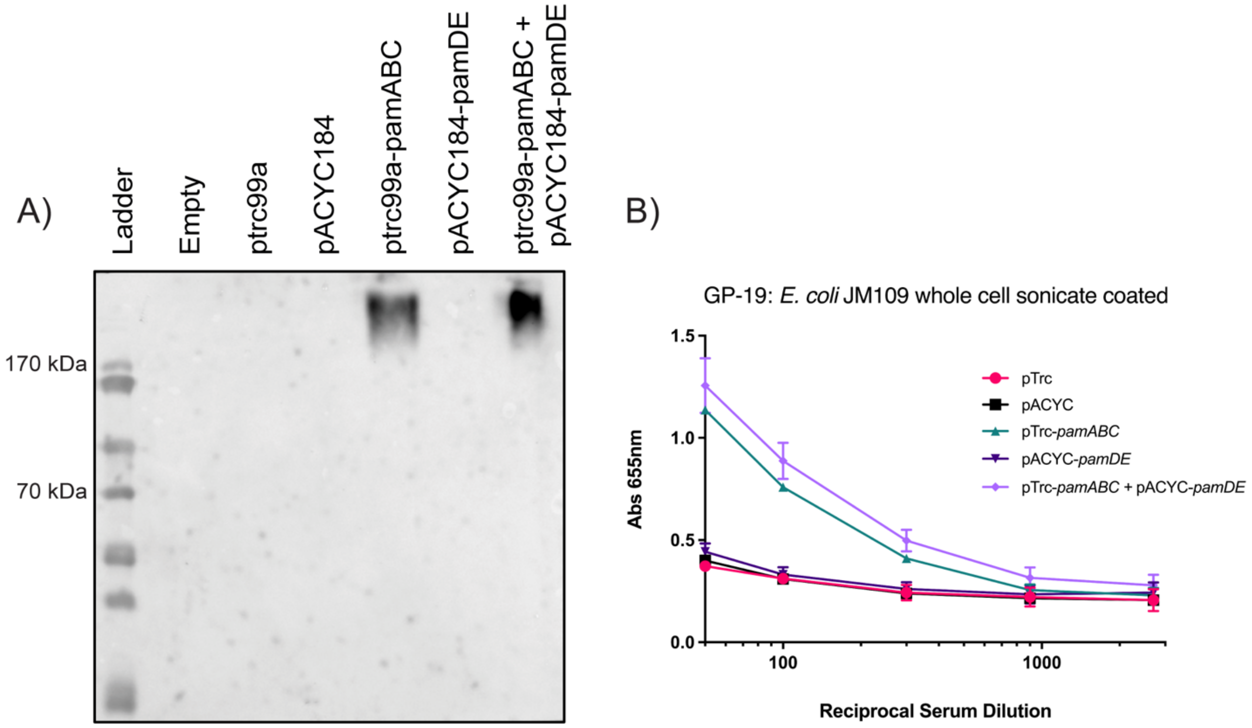


**Fig. S1: GP-19 antiserum is specific for the polymer synthesized by *pamABC* in *E. coli* JM109.** A) Galactan was detected by Western blot with GP-19 in whole cell sonicates from strains *E. coli* JM109 pTrc, JM109 pACYC, JM109 pTrc-*pamABC,* JM109 pACYC-*pamDE,* JM109 pTrc-*pamABC* pACYC-*pamDE.* Representative image shown. B) Galactan was detected in whole cell sonicates by ELISA with the GP-19 antiserum. Data are expressed as mean ± SEM from three independent experiments.

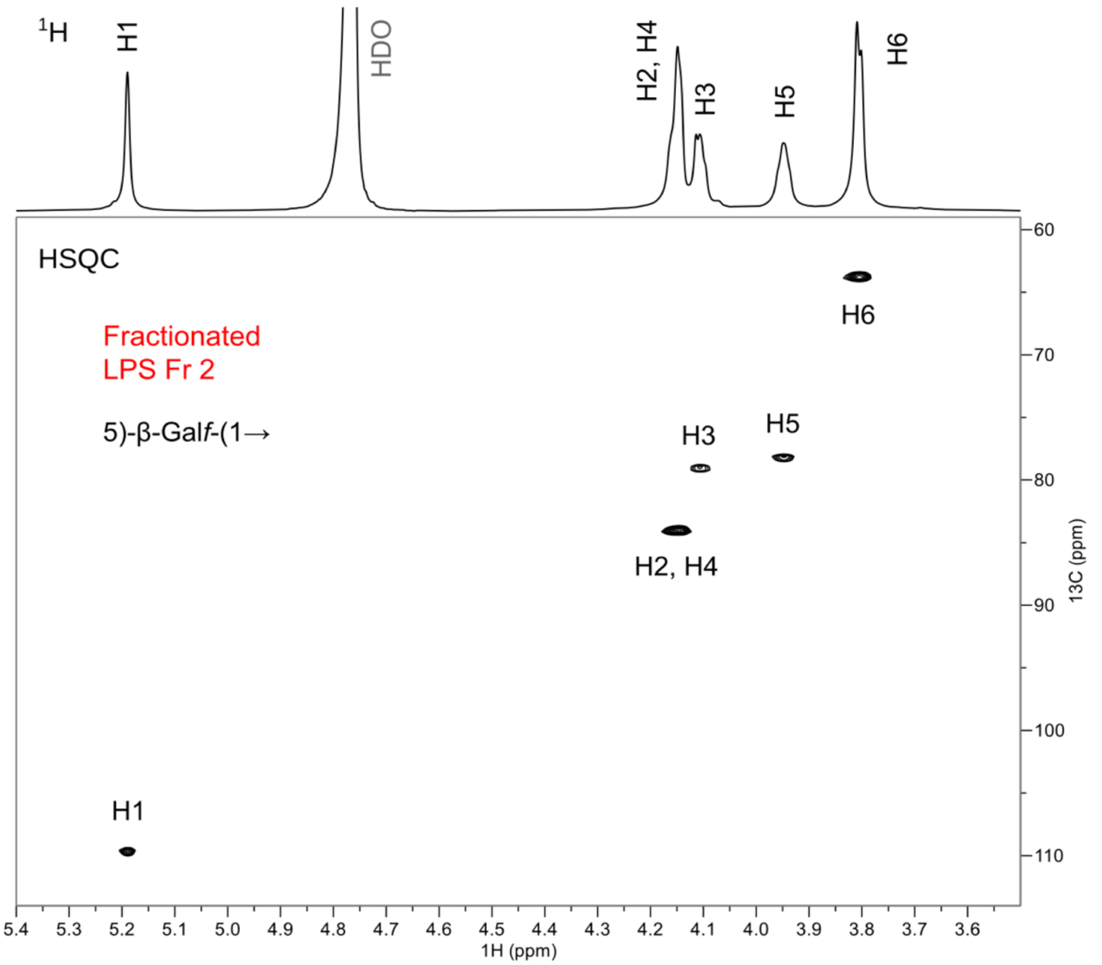


**Fig. S2: $^{1}$H and $^{1}$H,$^{13}$C-HSQC NMR spectra of LPS Fr. 2.** The HSQC signals of HMW LPS from Fraction 2 matched the galactan signals reported in Starr *et al*., 2013, as well as the signals of the galactan identified in the released carbohydrate material from Fr. 2 (Supplemental Fig. 3 and Supplemental Table 2).

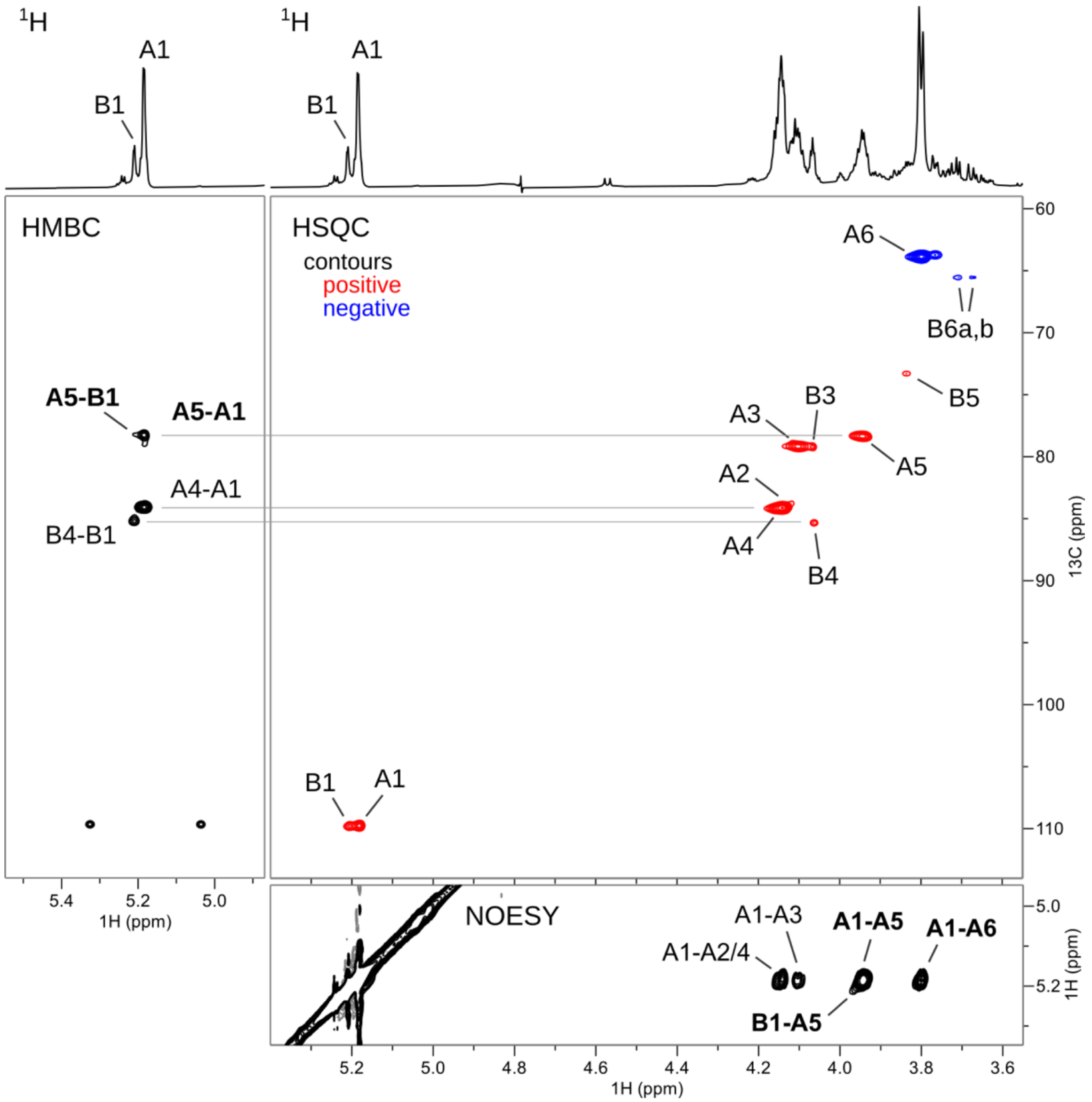


**Fig. S3: 2D NMR spectra of the carbohydrate material released from HMW LPS Fr. 2.** The HSQC signals of the carbohydrate released from HMW LPS Fr 2 were assigned with the aid of COSY, TOCSY, NOESY and HMBC NMR spectra. The partial NOESY and HMBC spectra are shown along the axes of the HSQC spectrum to illustrate signals instrumental in the assignment of the galactan resonances as well as identification of inter-residual correlations (marked in bold).

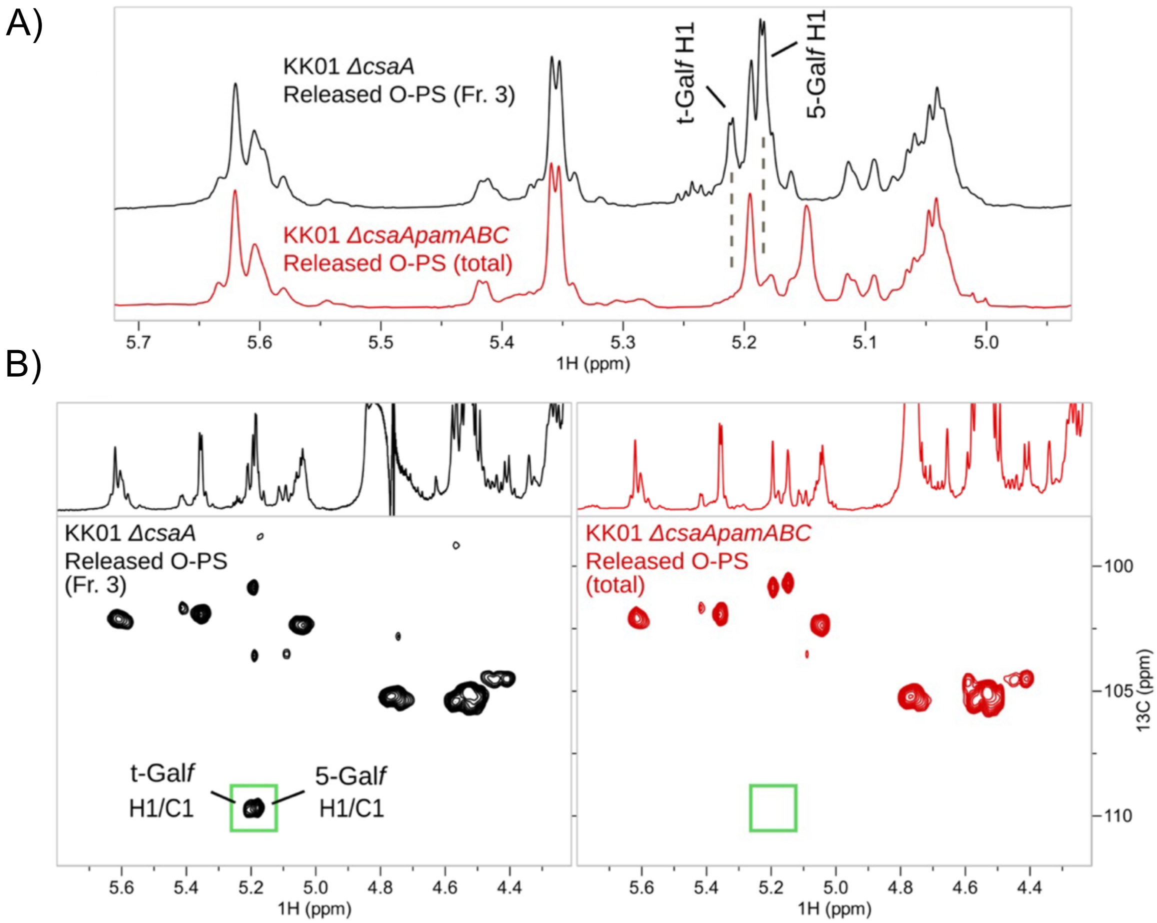


**Fig S4: Gal*f* is not present in the KK01 *ΔcsaApamABC* LPS.** A) Comparison of a partial anomeric region of $^{1}H$ NMR spectra of the KK01 *ΔcsaA* released LMW O-PS (from LMW LPS Fr. 3, in black) and the total, unfractionated O-PS released from KK01 *ΔcsaApamABC* LPS (in red). The signals of Gal*f* H1, present in the KK01 *ΔcsaA* O-PS, are absent from the KK01 *ΔcsaApamABC* O-PS. B) Comparison of the whole anomeric region of $^{1}H$,$^{13}C$-HSQC NMR spectra of the above O-PS material from the two mutants. Green rectangle highlights the positions of the anomeric Gal*f* signals that are not present in the KK01 *ΔcsaApamABC* O-PS spectrum. The corresponding regions of the $^{1}H$ spectra are shown above the HSQC spectra.

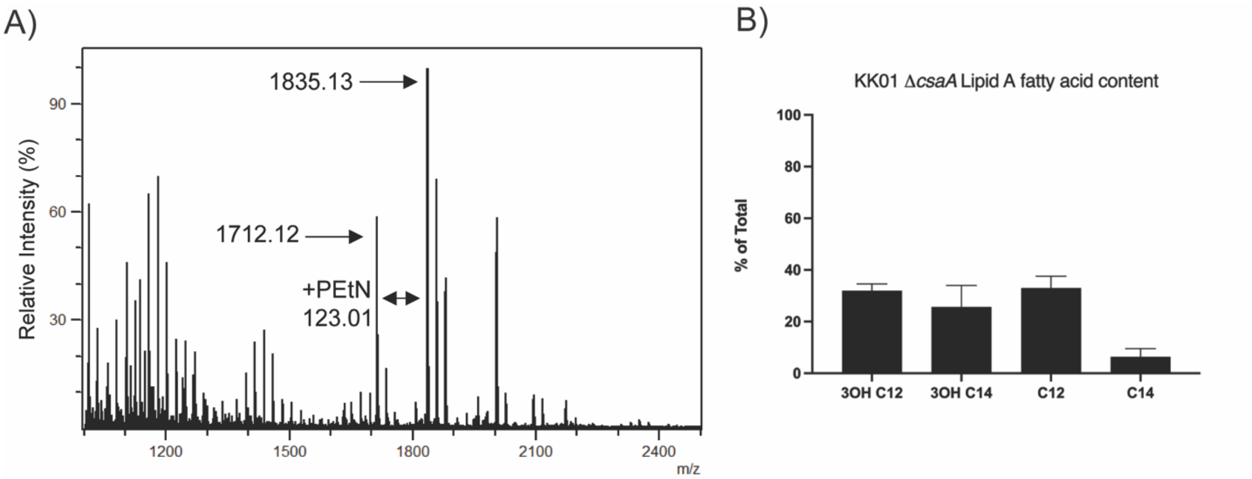


C) MS/MS of *m/z* 1712.12

| Naming | m/z (theory) | m/z (expt) | Error (Da) |
|---|---|---|---|
| $B_2$ | 1614.14 | 1614.08 | 0.06 |
| 3α | 1495.95 | 1495.89 | 0.06 |
| $B_2 + 3^{\alpha}$ | 1397.97 | 1397.92 | 0.05 |
| $B_2 + 3^{\alpha} + 3'\beta$ | 1199.81 | 1199.76 | 0.05 |
| $B_2 + 3\alpha + 2'\varepsilon$ | 1197.79 | 1197.75 | 0.04 |
| $B_2 + 3\alpha + 3'\beta + 2'\varepsilon$ | 999.63 | 999.59 | 0.04 |
| $B_2 + 3\alpha + 3'\alpha + 2'\varepsilon$ | 981.62 | 981.59 | 0.03 |
| $^{27}A_2 + 3'\alpha$ | 690.40 | 690.38 | 0.02 |

D) MS/MS of *m/z* 1835.13

| Naming | m/z (theory) | m/z (expt) | Error (Da) |
|---|---|---|---|
| $B'_2$ | 1737.15 | 1737.09 | 0.06 |
| 1 | 1712.12 | 1712.06 | 0.06 |
| $B'_2 + 1$ | 1614.15 | 1614.09 | 0.06 |
| $1 + B'_2 + 3'\alpha$ | 1397.97 | 1397.92 | 0.05 |
| $1 + B'_2 + 3'\alpha + 2'\varepsilon$ | 1197.79 | 1197.75 | 0.04 |
| $Y_1$ | 987.57 | 987.54 | 0.03 |
| $1 + {}^{27}A_2 + 3'\alpha$ | 690.40 | 690.38 | 0.02 |

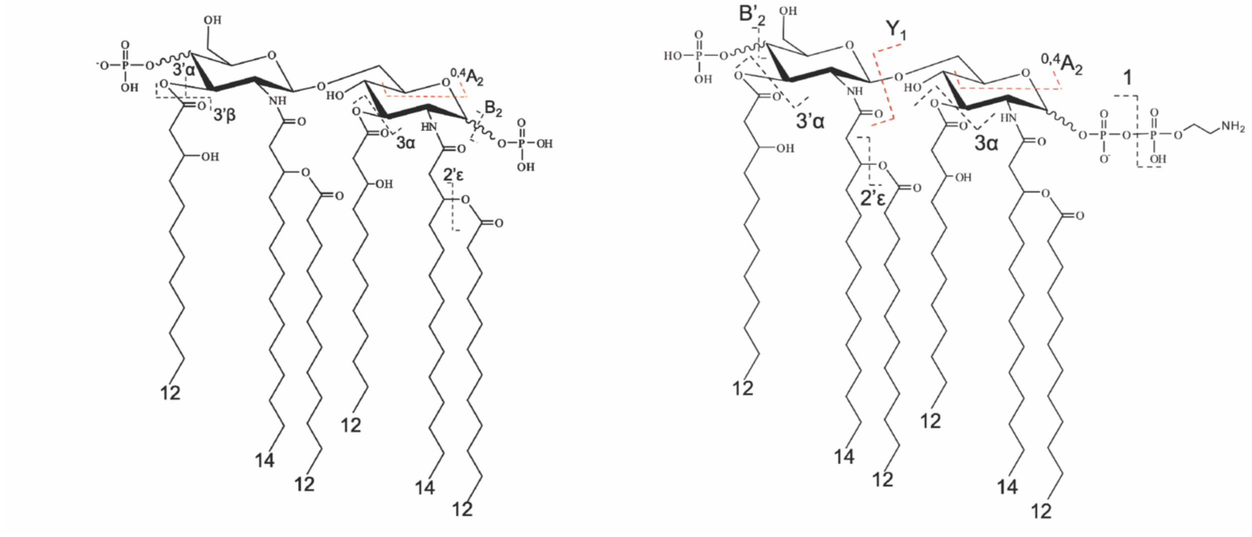

**Fig S5: Biochemical characterization of *K. kingae* lipid A.** A) MALDI mass spectrum obtained in negative ion mode from bacterial solution of *K. kingae* Δ*csaA* after FLAT processing. Major species detected as ions *m/z* 1712.12 and 1835.13. The *m/z* difference of 123.01 corresponds to addition of a phosphoethanolamine (PEtN) residue. B) *K. kingae* lipid A fatty acid content shown as a percentage of total. Lauric and myristic methyl esters are shown as 12:0 and 14:0, respectively. Hydroxylated methyl esters at the 3-position are denoted with 3-OH. C-D) Key fragment ions and corresponding structures identified by $FLAT^n$ and MS/MS for *K. kingae* lipid A species detected at ion *m/z* 1712.12 (C) and the PEtN modified lipid A detected at ion *m/z* 1835.13 (D).

**Table S1: Glycosyl composition of galactan prep for antibody production.**

| Residue | Mole % |
|---|---|
| Galactose (Gal) | 92.9 |
| Glucose (Glc) | 5.0 |
| N-acetyl Glucosamine (GlcNAc) | 2.1 |
| **Total** | **100.0** |

**Table S2: $^1$H and $^{13}$C chemical shift (ppm) assignments of galactan released from HMW LPS Fr. 2.**

| | Group | | | | | |
|---|---|---|---|---|---|---|
| Residue | **1** | **2** | **3** | **4** | **5** | **6(a,b)** |
| **A** | 5.19 | 4.14 | 4.11 | 4.16 | 3.95 | 3.80 |
| 5)-β-Gal*f*-(1→ | 109.6 | 84.0 | 79.0 | 84.1 | 78.2 | 63.7 |
| **B** | 5.22 | 4.15 | 4.07 | 4.07 | 3.84 | 3.72, 3.67 |
| t-β-Gal*f*-(1→ | 109.8 | 83.9 | 79.1 | 85.3 | 73.2 | 65.4 |

| Linkages | Method |
|---|---|
| A1 → A5 | NOESY, HMBC |
| B1 → A5 | NOESY, HMBC |

#### Table S3: Bacterial strains and plasmids used in this study.

| Strain or Plasmid | Description | Source or Reference |
|---|---|---|
| *K. kingae strains* | | |
| KK01 | Nonspreading/noncorroding derivative of clinical isolate 269–492 | [9] |
| KK01 Δ*csaA* | KK01 *csaA::un,* contains *csaA* deletion | This study |
| KK01 Δ*csaApamABCDE* | KK01 *csaA::un pam::kan,* contains *csaA* deletion and *pam* locus deletion | This study |
| KK01 Δ*csaApamABC* | KK01 *csaA::un pamABC::kan,* contains *csaA* deletion and *pamABC* deletion | This study |
| KK01 Δ*csaApamDE* | KK01 *csaA::un pamDE::kan,* contains *csaA* deletion and *pamDE* deletion | This study |
| KK01 Δ*csaApamABCrfaF* | KK01 *csaA::un pamABC::kan rfaF::erm,* contains *csaA* deletion, *pamABC* deletion and *rfaF* deletion | This study |
| *H. influenzae strains* | | |
| Rd | Non-adherent lab strain of *H. influenzae,* capsule-deficient serotype D | [10,11] |
| Rd Δ*rfaF* | Rd *rfaF::kan,* contains *rfaF* deletion | [11,12] |
| *E. coli strains* | | |
| DH5α | $\lambda^{-}$*ϕ*80d*lac*ZΔM15 Δ(*lacZYA-argF*)*U169 recA1 endA1* hsdR17($r_K^{-}$ $m_K^{-}$) *supE44 thi-1 gyrA relA1* | ThermoFisher Scientific |
| JM109 | *end*A1, *rec*A1, *gyr*A96, *thi, hsd*R17 ($r_k^{-}$, $m_k^{+}$), *rel*A1, *sup*E44, Δ( *lac-pro*AB), [F´ *tra*D36, *pro*AB, *laq*I$^{q}$ZΔM15]. | Promega |
| *Plasmids* | | |
| pTrc99a | High copy number expression vector containing the Trc promoter | [13] |
| pACYC184 | Low copy number plasmid vector containing the replication system of miniplasmid p15A | [14] |
| pUC19 | High copy number cloning vector | [15] |
| pFalcon2 | Source of the *aphA3* cassette | [16] |
| pIDN4 | Source of the *ermC* cassette | [17] |
| pUC19*csaA*::unmarked | Construct for introduction of an unmarked *csaA* deletion in *K. kingae* | [1] |
| pUC19*pam*::*ermC* | Construct for introduction of an *ermC*-marked *pamABCDE* deletion in *K. kingae* | [3] |
| pTrc99a-*pamABC* | pTrc99a containing *pamABC* under control of an IPTG-inducible promoter | This study |
| pACYC184-*pamDE* | pACYC184 containing *pamDE* under control of the native *pamD* promoter | This study |
| pΔ*pamABC* | pUC19-based plasmid for introduction of the *aphA3*-marked Δ*pamABC* deletion | This study |
| pΔ*pamDE* | pUC19-based plasmid for introduction of the *aphA3*-marked Δ*pamDE* deletion | This study |
| pΔ*rfaF* | pUC19-based plasmid for introduction of the *ermC*-marked Δ*rfaF* deletion | This study |

**Table S4: Primers used in this study**

| Primer Name | Sequence 5’ → 3’ |
|---|---|
| *pamA*5’_F | GCGAATTCGGCGTTGGTGGAATATCCTG |
| *pamA*5'_R | ACGTGGTACCACCTTCTGGTCGCTGAAATG |
| *pamC*3’_F | ACGTGGATCCCTATGTCGCAACGTTTGATAGG |
| *pamC*3’_R | ACGTAAGCTTGAATGTTGTCCAGCGCAATC |
| *aphA3*_F | GCATGGTACCCATCTAAATCTAGGTACTAAAACAATTCATCCAG |
| *aphA3*_R | GCATGGATCCGTTTGACAGCTTATCATCGATAAACCCAG |
| *pamD*5'_F | ACGTGAATTCTCGTGCTTATGCGCTGTTGC |
| *pamD*5’_R | ACGTGGTACCTTTCGGGATATTGCGGTTGG |
| *pamE*3’_F | GCGGATCCTCAAAGGCTGGTATAAACAC |
| *pamE*3’_R | GCAAGCTTCCATATCGCTTTGGCTTTGC |
| *rfaF5’*_F | ACGTGAATTCGGGAACAAGATTGCTGTAAC |
| *rfaF5’*_R | ACGTGGTACCCAATCCACGATGGGGAAATG |
| *rfaF3*’_F | ACGTGGATCCGCTTGCTGGACGAATTAAAC |
| *rfaF3’*_R | ACGTAAGCTTTAACGTGTCCCACAGGAATC |
| *ermC*_F | ACGTGGATCCGGTTACGCTTTGGGGAAATTATGAGG |
| *ermC*_R | ACGTGGTACCGTAATCATGGTCATAGCTGTTCGATAAGC |
| *pamABC*_F | AGCTGAATTCATGTTCCAATTAAGCGAAATTCC |
| *pamABC*_R | ACGTGGTACCTTTCGGGATATTGCGGTTGG |
| *pamDE*_F | GCGACCACACCCGTCCTGTGCTATGTCGCAACGTTTGATAG |
| *pamDE*_R | AAGGCTCTCAAGGGCATCGGACATTATTTAAATCCCAAATAATTCATAG |

#### SI References


1. Starr KF, Porsch EA, Seed PC, St Geme III JW. Genetic and Molecular Basis of Kingella kingae Encapsulation. Published online 2016. doi:10.1128/IAI.00128-16
2. Starr KF, Porsch EA, Seed PC, et al. Kingella kingae Expresses Four Structurally Distinct Polysaccharide Capsules That Differ in Their Correlation with Invasive Disease. Mitchell TJ, ed. *PLOS Pathog*. 2016;12(10):e1005944. doi:10.1371/journal.ppat.1005944
3. Starr KF, Porsch EA, Heiss C, Black I, Azadi P, St. Geme JW. Characterization of the Kingella kingae Polysaccharide Capsule and Exopolysaccharide. Adler B, ed. *PLoS One*. 2013;8(9):e75409. doi:10.1371/journal.pone.0075409
4. Manzi A, Esko J. Direct Chemical Analysis of Glycoconjugates for Carbohydrates. *Curr Protoc Mol Biol*. 1995;32(1):1-11. doi:10.1002/0471142727.mb1709s32
5. Lees A, Nelson BL, Mond JJ. Activation of soluble polysaccharides with 1-cyano-4-dimethylaminopyridinium tetrafluoroborate for use in protein-polysaccharide conjugate vaccines and immunological reagents. *Vaccine*. 1996;14(3):190-198. doi:10.1016/0264-410X(95)00195-7
6. Yang H, Jackson SN, Woods AS, Goodlett DR, Ernst RK, Scott AJ. Streamlined Analysis of Cardiolipins in Prokaryotic and Eukaryotic Samples Using a

Norharmane Matrix by MALDI-MSI. *J Am Soc Mass Spectrom*. 2020;31(12):2495-2502. doi:10.1021/jasms.0c00201

7. Niedermeyer THJ, Strohalm M. mMass as a Software Tool for the Annotation of Cyclic Peptide Tandem Mass Spectra. *PLoS One*. 2012;7(9). doi:10.1371/journal.pone.0044913
8. Wollenweber HW, Rietschel ET. Analysis of lipopolysaccharide (lipid A) fatty acids. *J Microbiol Methods*. 1990;11(3-4):195-211. doi:10.1016/0167-7012(90)90056-C
9. Kehl-Fie TE, St. Geme JW. Identification and characterization of an RTX toxin in the emerging pathogen Kingella kingae. *J Bacteriol*. 2007;189(2):430-436. doi:10.1128/JB.01319-06
10. Setlow JK, Randolph ML, Boling ME, Mattingly A, Price G, Gordon MP. Repair of DNA in Haemophilus Influenzae. II. Excision, Repair of Single-Strand Breaks, Defects in Transformation, and Host Cell Modification in UV-Sensitive Mutants. *Cold Spring Harb Symp Quant Biol*. 1968;33:209-218. doi:10.1101/SQB.1968.033.01.024
11. Spahich NA, Hood DW, Moxon ER, St Geme JW. Inactivation of Haemophilus influenzae lipopolysaccharide biosynthesis genes interferes with outer membrane localization of the hap autotransporter. *J Bacteriol*. 2012;194(7):1815-1822. doi:10.1128/JB.06316-11
12. Nichols WA, Gibson BW, Melaugh W, Lee NAG, Sunshine M, Apicella MA. Identification of the ADP-L-glycero-D-manno-heptose-6-epimerase (rfaD) and heptosyltransferase II (rfaF) biosynthesis genes from nontypeable Haemophilus influenzae 2019. *Infect Immun*. 1997;65(4):1377. doi:10.1128/iai.65.4.1377-1386.1997
13. Amann E, Ochs B, Abel KJ. Tightly regulated tac promoter vectors useful for the expression of unfused and fused proteins in Escherichia coli. *Gene*. 1988;69(2):301-315. doi:10.1016/0378-1119(88)90440-4
14. Chang ACY, Cohen SN. Construction and Characterization of Amplifiable Multicopy DNA Cloning Vehicles Derived from the P15A Cryptic Miniplasmid. 1978;134(3):1141-1156.
15. Yanisch-Perron C, Vieira J, Messing J. Improved M13 phage cloning vectors and host strains: nucleotide sequences of the M13mpl8 and pUC19 vectors. *Gene*. 1985;33(1):103-119. doi:10.1016/0378-1119(85)90120-9
16. Hendrixson DR, Akerley BJ, DiRita VJ. Transposon mutagenesis of Campylobacter jejuni identifies a bipartite energy taxis system required for motility. *Mol Microbiol*. 2001;40(1):214-224. doi:10.1046/J.1365-2958.2001.02376.X
17. Hamilton HL, Schwartz KJ, Dillard JP. Insertion-Duplication Mutagenesis of Neisseria: Use in Characterization of DNA Transfer Genes in the Gonococcal Genetic Island. *J Bacteriol*. 2001;183(16):4718. doi:10.1128/JB.183.16.4718-4726.2001
